# Small molecule agonists of 8-oxoguanine DNA glycosylase, OGG1

**DOI:** 10.64898/2026.01.30.702659

**Authors:** Michael M. Luzadder, Irina G. Minko, Samantha A. Moellmer-Gomez, Naoto N. Tozaki, Pawel Jaruga, Miral Dizdaroglu, Haihong Jin, Jordan Devereaux, Aaron Nilsen, R. Stephen Lloyd, Amanda K. McCullough

**Affiliations:** Oregon Institute of Occupational Health Sciences, Oregon Health & Science University, Portland, OR 97239; Biomolecular Measurement Division, National Institute of Standards and Technology, Gaithersburg, MD, 20899; Department of Chemical Physiology & Biochemistry, Oregon Health & Science University, Portland, OR 97239; Department of Molecular and Medical Genetics, Oregon Health & Science University, Portland, OR 97239

**Author notes:** Contributed Equally (Michael M. Luzadder, Irina G. Minko). Department of Molecular, Cellular, and Developmental Biology, and the Interdisciplinary Quantitative Biology Program (IQ Biology), BioFrontiers Institute, University of Colorado, Boulder, CO 80309, USA. Department of Biomedical Engineering, Oregon Health & Science University, Portland, OR 97239. Abbott Core Diagnostics, Abbott Park, IL 60064. For correspondence: Amanda K. McCullough,; R. Stephen Lloyd.

**Keywords:** DNA repair, reactive oxygen species, DNA glycosylase, apurinic/apyrimidinic lyase, paraquat

## Abstract

Base excision repair (BER) is the primary pathway that removes oxidatively-induced DNA base damage from the nuclear and mitochondrial genomes, with 8-oxoguanine DNA glycosylase (OGG1) initiating repair at the two most frequently-formed base lesions: 8-oxo-7,8-dihydro-2′-deoxyguanosine (8-oxoGua) and 2,6-diamino-4-oxo-5-formamidopyrimidine (FapyGua). Humans expressing a catalytically-compromised variant of OGG1 (S326C) are at increased risk for type 2 diabetes, Alzheimer’s disease, and Parkinson’s disease. To potentially enhance the overall catalytic efficiency of this variant, a prior medicinal chemistry screen discovered seven chemically distinct agonists of OGG1 that stimulated activity *in vitro* and attenuated a paraquat (PQ) challenge in cultured cells. Herein, we developed structure-activity relationships around one specific core structure, F01. Using fluorescence-based DNA cleavage assays, we assessed the abilities of these compounds to stimulate the overall rate of OGG1 catalysis. Multiple compounds were identified that increased OGG1 activity on DNAs containing a site-specific 8-oxoGua by 2-fold or greater, with 9 compounds showing EC_50_ concentrations lower than F01 and were specific for OGG1. Selected agonists were shown to enhance OGG1-catalyzed release of 8-oxoGua and FapyGua from γ-irradiated high-molecular-weight DNA using gas chromatography tandem mass spectrometry analyses. Since these assays did not reveal which step in the overall reaction was stimulated, we used a separation-of-function OGG1 mutant that possessed glycosylase, but not abasic-site (AP) lyase activity to demonstrate that the glycosylase step was not enhanced. In contrast, all agonists stimulated the AP lyase activity to levels equal to or greater than the magnitude of stimulation observed for overall chemistry, implicating agonist-mediated turnover as a potential contributor to the overall rate stimulation. The biological activities of selected agonists were evaluated in OGG1-deficient Kasumi-1 cells under conditions of paraquat (PQ)-induced oxidative stress, with several compounds mitigating PQ challenge.

## Introduction

Reactive oxygen species (ROS), produced by both normal cellular metabolism and exogenous sources, create oxidatively-induced DNA damage in both the nuclear and mitochondrial genomes (1, 2). Mitochondrial DNA is especially vulnerable to this type of damage due to its proximity to ROS production in the inner mitochondrial membrane. Oxidatively-induced base lesions are primarily repaired by the base excision repair (BER) pathway in which both damage recognition and hydrolysis of the N-glycosidic bond between the deoxyribose and the damaged base are carried out by DNA glycosylases that in humans include OGG1, NTHL1, NEIL1, NEIL2, and NEIL3 (3). A subset of these lesions, 8-oxo-7,8-dihydro-2′-deoxyguanosine (8-oxoGua) and 2,6-diamino-4-oxo-5-formamidopyrimidine (FapyGua) are removed with comparable efficiencies by OGG1, producing an apurinic/apyrimidinic (AP) site (4–7). This AP site is further processed by either OGG1 which can catalyze a β-elimination reaction producing a 3′-phospho α,β-unsaturated aldehyde (dRP) and a 5′ phosphate or AP endonuclease 1 (APE1) generating a 3′-OH and a 5′ dRP (8, 9). Prior investigations have suggested that OGG1 dissociation from the AP site constitutes the rate-limiting step in the overall reaction (10). Once an appropriate 3′-OH is generated, repair is completed by the sequential activities of DNA polymerases and ligases. However, if DNA replication occurs prior to repair, 8-oxoGua and FapyGua lesions are potentially mutagenic, giving rise to G to T transversions and G to A transitions (11–15).

Epidemiological studies have connected OGG1 deficiency to a variety of human diseases (9) including Alzheimer’s (16, 17) and Parkinson’s diseases (PD) (18), obesity (19), type 2 diabetes (20), and cardiovascular disease (21). Deficiencies in OGG1 are also associated with mitochondrial dysfunction and ATP depletion (22, 23). Additionally, elevated levels of 8-oxoGua lesions have been found in the mitochondria of dopaminergic neurons in PD brains and have been associated with dopaminergic neuronal loss (24). The S326C OGG1 variant has reduced catalytic efficiency *in vitro* and shows positive correlations with a variety of cancers (25–30).

In murine knock-out models, *Ogg1^-^*^/-^ mice develop age- and diet-induced metabolic syndrome characterized by insulinemia, fatty liver, adipose tissue inflammation, and obesity (31). Additionally, *Ogg1^-^*^/-^mice show decreased muscle function and changes in microbiome composition (32, 33). In contrast, transgenic mice that overexpress human mitochondrially-targeted OGG1 (mtOGG1) are resistant to the development of diet-induced metabolic syndrome, show increased tissue mitochondrial content, and decreased levels of inflammation (34–36). Additionally, yellow agouti mice, characterized by genetically-driven metabolic syndrome, similarly benefit from the overexpression of human mitochondrially-targeted OGG1 (34). These outcomes are achieved when human mtOGG1 is overexpressed at levels only 2-3-fold higher than mtOGG1 levels in WT mice. Collectively, these observations suggest benefits of enhanced mtOGG1 activity and position OGG1 as a target for small molecule activation.

To address this goal of enhancing OGG1 catalytic efficiency, there are several steps along the reaction coordinate that could potentially achieve this objective. Comparable to other DNA glycosylases, the full catalytic cycle of OGG1 involves nonspecific DNA binding and translocation, specific base damage recognition involving nucleotide flipping into an active-site pocket, sequential DNA glycosylase and AP lyase reactions, followed by enzyme-product dissociation. Thus, stimulation of the glycosylase, or AP lyase chemistries or promotion of enzyme dissociation could prove beneficial.

The Verdine laboratory was the first to report stimulation of BER by small molecule guanine analogs: 8-bromoguanine (8-bromoGua) and 8-aminoguanine (8-aminoGua) (37). These were hypothesized to stimulate OGG1 activity through a mechanism of product-assisted catalysis in which 8-bromoGua and 8-aminoGua were bound in the product-recognition pocket of OGG1. However, recent investigations favor an allosteric mechanism of OGG1 activation in which guanine analogs bind outside of the product recognition pocket (38). Allosteric activation of OGG1 by guanine analogs is reported to increase the rate of the OGG1-catalyzed AP-lyase reaction (38).

Since this initial report of OGG1 activation, other groups have used screening approaches designed to identify other small molecule agonists of OGG1 ((39, 40); and reviewed in (41)). Baptiste et al. (39) used an *in vitro* fluorescence-based DNA cleavage assay to identify several compounds that increased the overall reaction rate of OGG1. These compounds were shown to enhance the removal of 8-oxoGua in the mitochondrial genome and improve mitochondrial function under conditions of oxidative stress induced by PQ (39). Expanding on these findings, Michel et al. (40) further characterized compound C (TH10785) previously identified by Baptiste et al (39). Mechanistically, this molecule confers OGG1 with the novel ability to catalyze δ-elimination reactions at AP sites, in addition to its intrinsic ability to catalyze β-elimination. Cocrystal structures of OGG1 and TH10785 indicate binding of the ligand in the active site pocket of OGG1 (40). Recent investigations have reported on organocatalytic switches to enhance the AP lyase activity of OGG1 (42).

The objective of our study was to identify and characterize new small molecule agonists of OGG1. Our experimental strategy focused on developing a medicinal chemistry-based, structure-activity relationship primarily using a previously described potent agonist of OGG1, 1-cyclohexyl−1-(2,4-dichlorophenyl)−2-(1H-imidazol−1-yl) ethanol (39), (F01; Fig. 1A) as a common core structure and other closely related derivatives. To identify additional OGG1 agonists and to gain insights into which step in the overall reaction coordinate was stimulated, our strategy utilized a variety of techniques that measured either i) an overall stimulation of OGG1 activity at any step in the reaction, or ii) stimulation of the AP lyase activity alone; or iii) stimulation of only the glycosylase activity; or iv) stimulation of either the glycosylase activity or enzyme turnover.

**Figure 1.**
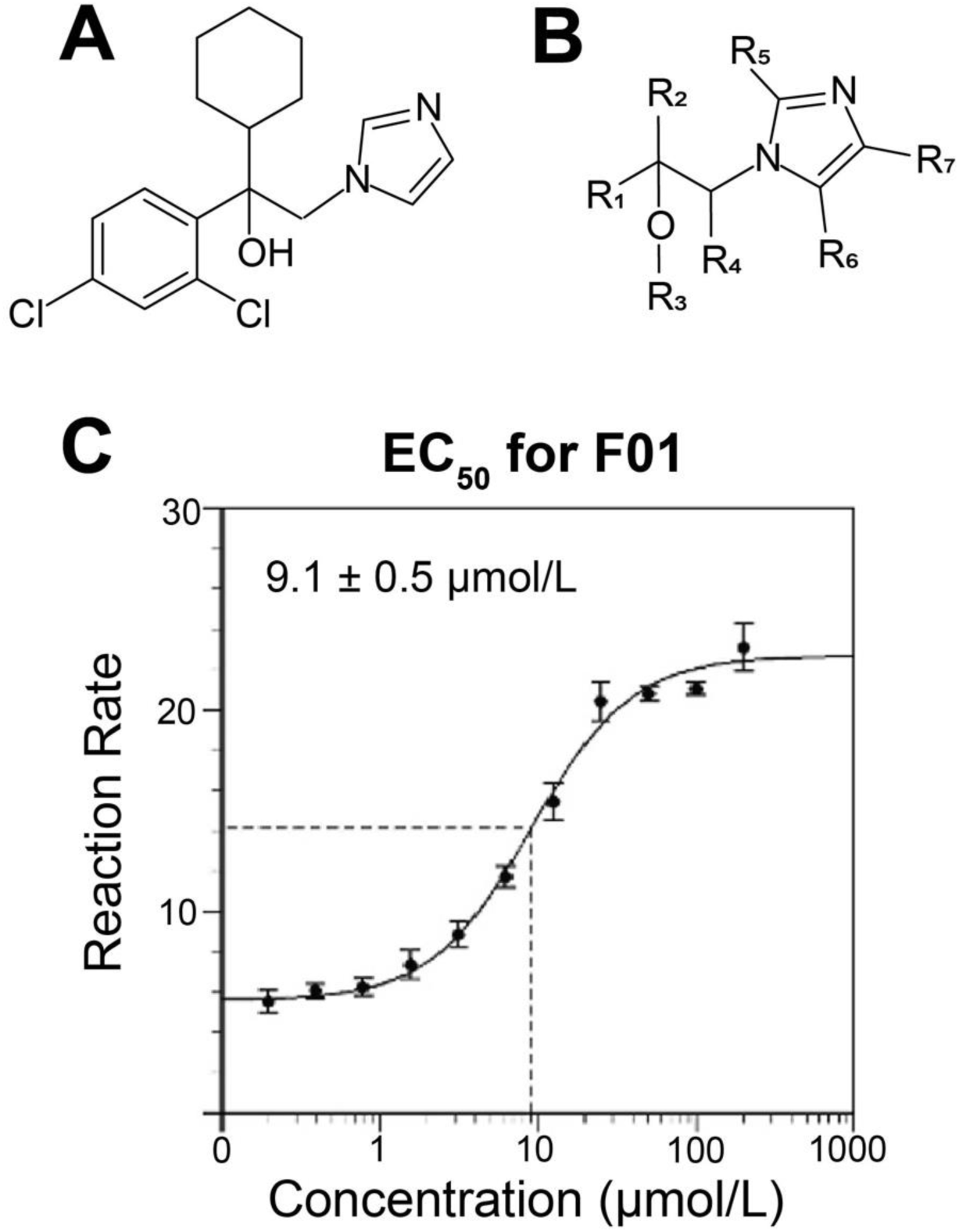
*A*, 1-cyclohexyl− 1-(2,4-dichlorophenyl)− 2-(1H-imidazol−1-yl) ethanol (F01), OGG1 agonist reported by Baptiste et al. in 2018 (39). *B*, Core molecular structure of OGG1 agonist compounds with specified substitution positions. *C,* EC_50_ for F01 with 8-oxoGua-containing DNA substrate. Uncertainties are standard deviations.

### Experimental procedures

#### Materials

OGG1 agonist candidate compounds were purchased from Molport (Riga, Latvia; https://www.molport.com) and ChemBridge (San Diego, CA, USA; https://chembridge.com). TH10785 was purchased from MedChemExpress (catalog # HY-147131) and TargetMol (catalog number T60034). Compounds F50 and F51 were purchased from Toronto Research Chemicals (catalog # TRC-E326000 and # TRC-M342490), respectively. F51 was also purchased from US Biological Life Sciences (catalog # 164979).

The source identification number, International Union of Pure and Applied Chemistry (IUPAC) name, molecular formula and molecular weight for each compound can be found in Table S1. All compounds were reconstituted in 100 % DMSO at 10 mmol/L.

The 5’-TAMRA-labeled oligodeoxynucleotide containing an internal 8-oxoGua was synthesized as previously described (43). All other oligodeoxynucleotides were synthesized by Integrated DNA Technologies (Coralville, IA). Uracil DNA Glycosylase (UDG) was purchased from New England Biolabs (Ipswich, MA, USA; RRID:SCR_013517). AlamarBlue was purchased from Bio-Rad (Hercules, CA, USA; RRID:SCR_008426) (catalog #: BUFO12B). Paraquat dichloride hydrate (1,1’-dimethyl-4,4’-bipyridinium dichloride) was purchased from Sigma-Aldrich (St. Louis, MO, USA; RRID:SCR_008988) (catalog #: 75365-73-0). PQ was reconstituted in deionized water at 50 mmol/L.

#### Purification of DNA glycosylases

Human OGG1 and the separation-of-function mutant KCCK were expressed in *Escherichia coli* (*E. coli*) using plasmid vectors that introduced a N-terminal 6-His affinity purification tag. These enzymes were purified as previously published (44). Wild-type (WT) edited Nei-like DNA glycosylase 1 (NEIL1) was purified as previously described (45). *E. coli* formamidopyrimidine DNA glycosylase (Fpg) was purified as previously described (46). An expression vector for human endonuclease III-like protein 1 (NTH1) was kindly provided by Dr. Susan Wallace (University of Vermont, Burlington, VT). NTH1 was purified using Ni-NTA affinity chromatography using methods described for NEIL1 (45).

#### DNA substrates

The oligodeoxynucleotides used for DNA cleavage assays in this study included: (1) 5’-TAMRA-labeled 17-mer containing an internal 8-oxoGua (5’-TAMRA-TCACC(8-oxoGua)TCGTACGACTC-3’); (2) 5’-TAMRA-labeled 17-mer containing an internal uracil (U) (5’-TAMRA-TCACCUTCGTACGACTC-3’); (3) 5’-TAMRA-labeled 17-mer containing an internal thymine glycol (ThyGly) (5’-TAMRA-TCACCT(ThyGly)CGTACGACTC-3’); (4) a complementary 17-mer conjugated with BHQ2 at its 3’ terminus and a cytosine opposite 8-oxoGua or U or an adenine opposite ThyGly. To prepare double-stranded DNA substrates, 1 µmol/L 5’-TAMRA conjugated 8-oxoGua-, U-, or ThyGly-containing oligodeoxynucleotides were combined with 1.2 µmol/L BHQ2-conjugated complementary oligodeoxynucleotides in 20 mmol/L Tris-HCl buffer (pH 7.4) containing 100 mmol/L KCl and 0.01 % (v/v) Tween-20 and heated to 90 °C for 2 min then slowly cooled to 4 °C. AP sites were produced from U-containing oligodeoxynucleotides by incubation of 100 nmol/L DNA with UDG (0.5 U/µL) for 30 min at 37 °C in 20 mmol/L Tris-acetate (pH 7.9), 50 mmol/L potassium acetate, 10 mmol/L magnesium acetate, and 100 µg/mL BSA (CutSmart buffer from New England Biolabs Inc.).

#### Fluorescence-based DNA cleavage assay

Screening reactions were performed at 37 °C in a reaction volume of 20 µL. WT OGG1 was diluted to 125 nmol/L in buffer containing 20 mmol/L Tris-HCl (pH 7.4), 100 mmol/L KCl, 0.01 % Tween-20, and 100 µg/mL bovine serum albumin (BSA). Test compounds were diluted to 100 µmol/L in 100 % DMSO prior to screening. For each reaction, 32 µL of 125 nmol/L WT OGG1 was pre-mixed with 8 µL of 100 µmol/L test compound. Reactions were initiated by the addition of 10 µL of the OGG1 + compound mixture to 10 µL of 100 nmol/L AP site- or 8-oxoGua-containing oligodeoxynucleotides in a 384 well black flat-bottom microplate using a multichannel pipette. Reaction plates were centrifuged for 1 min at 200 x g prior to measurement at 37 °C in a TECAN INFINITE M NANO instrument. The final concentrations of reactants in all screening reactions were as follows: 50 nmol/L WT OGG1, 50 nmol/L 8-oxoGua- or AP-site-containing oligodeoxynucleotide, 10 µmol/L test compound and 10 % (v/v) DMSO. Fluorescence readings were taken every 2 min for 1 h using a 525 nm (9 nm bandwidth) excitation filter and a 598 nm (20 nm bandwidth) emission filter. All reactions were performed in technical duplicate. Readings from technical duplicates were averaged, and the initial rate was determined by fitting the data of the linear phase of the reaction to a linear function using Excel. The initial rate was divided by that of the OGG1 + DMSO control reaction to calculate the relative fold change. Agonists (10 µmol/L) were also screened with WT edited NEIL1 (15 nmol/L) with ThyGly-containing oligodeoxynucleotide (50 nmol/L) and Fpg (15 nmol/L) with 8-oxoGua-containing oligodeoxynucleotide under the same assay conditions described above.

#### EC_50_ measurements

EC_50_ curves were generated for selected agonists using 9 to14 agonist concentrations with 8-oxoGua-containing oligodeoxynucleotide. OGG1 agonists were diluted to 10-fold of the final concentration in 100 % DMSO. Reactions were performed and initial rates calculated as described above for screening. The rates for each agonist concentration were plotted, and non-linear regressions were performed to determine the EC_50_ concentrations using the ATT Bioquest EC_50_ calculator (47). The data were fitted to the following equation using four parameter mode. 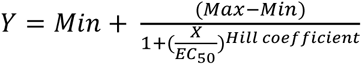

Each agonist was tested in three independent experiments, and the mean EC_50_ was calculated with the standard deviation shown.

#### Measurements of oxidatively-induced DNA bases by gas chromatography-tandem mass spectrometry

Triplicates of dried DNA samples (50 µg each) were supplemented with aliquots of stable isotope-labeled analogues of modified DNA bases as internal standards, incubated with 2 µg of OGG1 for 10 min to release modified DNA bases from DNA, and then analyzed by GC-MS/MS as described (48). Agonists were tested at 2x EC_50_ concentrations.

#### Gel-based DNA cleavage assays with KCCK OGG1 mutant

Gel-based DNA cleavage assays were performed using the KCCK OGG1 mutant and selected agonists with 8-oxoGua-containing oligodeoxynucleotide. KCCK OGG1 was diluted to 62.5 nmol/L in buffer containing 20 mmol/L Tris-HCl (pH 7.4), 100 mmol/L KCl, 0.01 % Tween-20, and 100 µg/mL BSA. Test compounds were diluted to 100 mmol/L in 100 % DMSO. For each reaction, 32 µl of 62.5 nmol/L KCCK OGG1 were premixed with 8 µl of 100 µmol/L agonist or 100 % DMSO. Reactions were initiated by the addition of 10 µL of 200 nmol/L 8-oxoGua-containing oligodeoxynucleotide substrate to 10 µL of each KCCK OGG1 mutant and agonist mixture. Reactions were performed in a 37 °C manifold heat block. After 15 min, reactions were terminated with the addition of 20 µL of 0.5 mol/L NaOH and incubated at 90 °C for 2 min. Two volumes of formamide solution with 10 mmol/L EDTA were added to each reaction. DNAs were resolved by electrophoresis through a 15 % polyacrylamide gel in the presence of 8 mol/L urea. TAMRA-conjugated DNAs were visualized using a FluorChem M imager (Protein Simple) with a 534 nm LED light source and 593 nm emission filter. The intensities of TAMRA fluorescence that were associated with the DNA bands were measured by the FluorChem M built-in software. Data were analyzed and plotted using GraphPad Prism software.

#### Cell line

The human leukemia cell line Kasumi-1 was generously provided by Dr. Jeffery Tyner, OHSU. The Kasumi-1 cell line is characterized by a RUNX1-RUNX1T1 fusion and reduced levels of nuclear OGG1 (49). Kasumi-1 cells were cultured under sterile conditions in T25 and T75 flasks (Greiner Bio-One) using RPMI 1640 culture media (HyClone) supplemented with 20 % fetal bovine serum (FBS; Corning). Kasumi-1 cells were maintained in suspension culture at a density of 0.5 x 10^6^ cells/mL to 1.5 x 10^6^ cells/mL in a humidified ambient oxygen incubator at 37 °C with 5 % CO_2_. Cell counts were taken using a hemocytometer and trypan blue exclusion. Kasumi-1 cells were routinely tested for mycoplasma contamination, and the cells were authenticated as previously described (49).

#### Cytotoxicity assays

Exponentially-growing Kasumi-1 cells were seeded in 96 well plates at ≍10000 cells per well in 90 µL of RPMI-1640 media and incubated overnight at 37 °C and 5 % CO_2_. Cells were treated with increasing concentrations of PQ (0 µmol/L to 500 µmol/L) by addition of 10 µL of PQ diluted in culture media. All treatments were performed in technical triplicate, in which three identical wells were plated for each condition tested. After 72 h at 37 °C and 5 % CO_2_, metabolic activity was assayed by adding 10 µL of AlamarBlue to each well and incubating cells for 5 h. Fluorescence readings were taken on a TECAN INFINITE M NANO plate reader using a 545 nm (9 nm bandwidth) excitation filter and a 590 nm (20 nm bandwidth) emission filter. Fluorescence readings from each well and the technical triplicates were averaged, and the background fluorescence of culture media alone was subtracted. The experimental wells containing PQ-treated cells were normalized to untreated control cells. To establish whether treatment with agonists alone affected cellular metabolism, Kasumi cells were also treated with increasing concentrations of agonist alone (0 µmol/L to 10 µmol/L) and percent metabolic activity was measured as described above.

For combination treatments of PQ and agonists, exponentially growing Kasumi-1 cells were seeded as described above. Cells were treated with agonists and 3 h later with PQ. Treatments were performed by adding 10 µL of agonist compounds, diluted in culture media to 100 µmol/L, to each well in technical triplicate. DMSO was held constant at a final concentration of 0.1 % (v/v) between agonist concentrations. After 3 h, cells were treated with PQ at a final concentration of 250 µmol/L. After 72 h at 37 °C and 5 % CO_2_, metabolic activity was assayed by AlamarBlue with fluorescence readings taken as described above. Fluorescence readings from experimental wells containing treated cells were then normalized to cells treated with only 0.1 % DMSO. All cytotoxicity assays were performed at a minimum, three independent times. Data were analyzed and plotted using Graph Pad Prism software.

## Results

### Experimental rationale

Based on the initial biochemical and cellular characterizations of the efficacy of F01 (1-cyclohexyl− 1-(2,4-dichlorophenyl)− 2-(1H-imidazol−1-yl)ethanol) (Fig. 1A) as a potent and biologically effective agonist of OGG1 (39), we hypothesized that the potency of this molecule could be increased via structural optimization. To address this hypothesis, since F01 was not commercially available, it was synthesized as described in the Materials and Methods section and schematically shown in Fig. S1. We then assembled an additional library of analogs of F01 (Table S1). Of these, 52 compounds were commercially available, and 5 compounds (F55, F56, F57, F58, and F59) were synthesized as part of this investigation. The structures of 51 of the 59 compounds were generalized to a common core structure with specified substitution positions R_1_ - R_7_ (Fig. 1B). Based on R_1_ substitutions to the core structure, 51 compounds were classified into six chemotypes: phenyl (Table 1), 4-OH-phenyl (Table 2), halogenated phenyl (Table 3), 2,4-dichloro phenyl (Table 4), CH_2_-O-phenyl (Table 5), and pyridyl (Table 6). Eight other compounds that did not share a common core structure were classified separately (Table 7). All compounds were screened for stimulatory effects on the overall OGG1 activity and evaluated using various DNA cleavage assays to further characterize their activity and mechanism. Additionally, the biological activity of selected agonists was evaluated under conditions of PQ-induced oxidative stress using cellular metabolic activity as an endpoint.

**Table 1.**
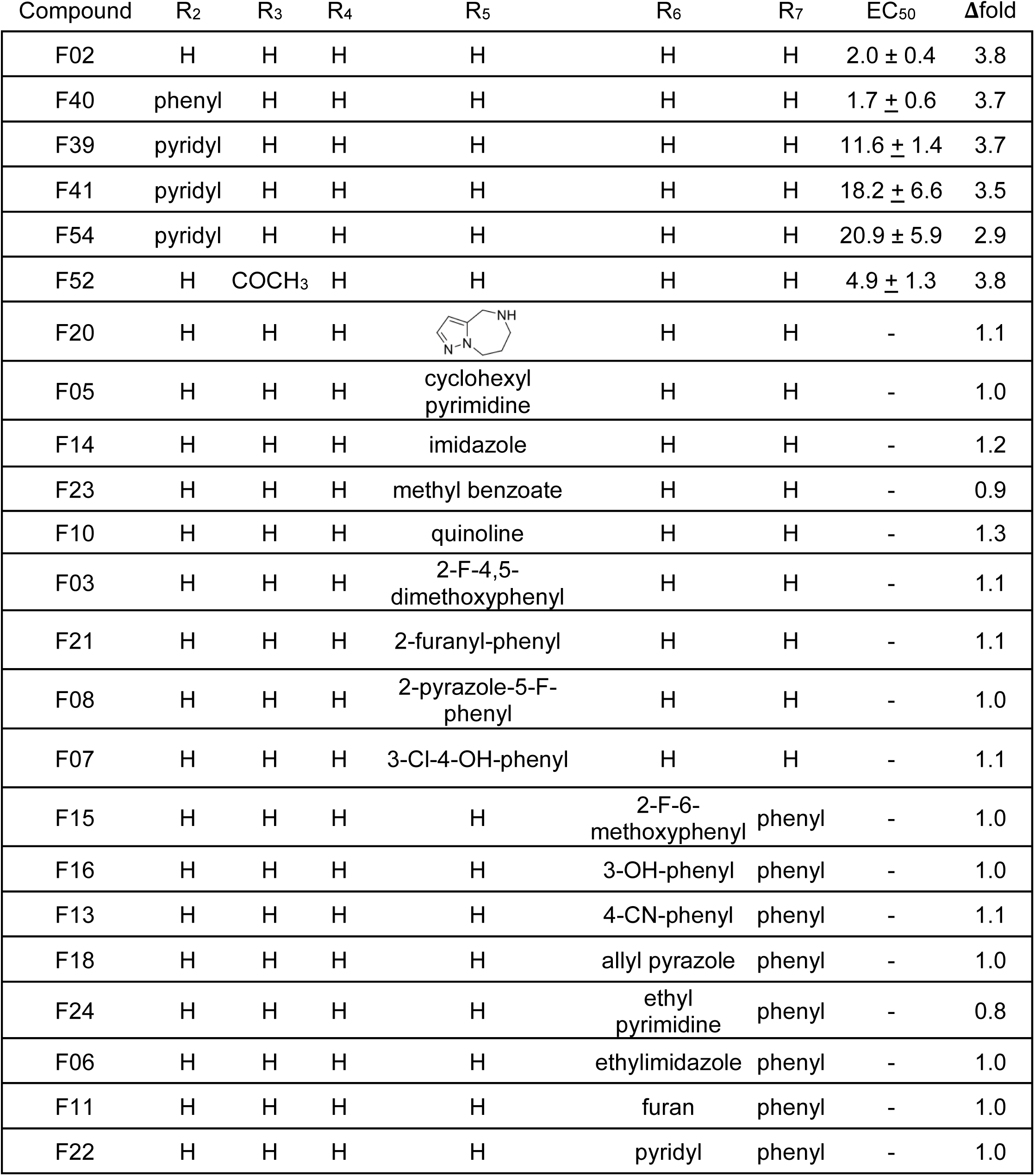
R_1_ phenyls. The fold change of the initial reaction rate of OGG1 (50 nmol/L) with 8-oxoGua-containing DNA substrate (50 nmol/L) in the presence of agonists (10 µmol/L) is given relative to the DMSO control. EC_50_ concentrations determined by analyzing initial reaction rates of OGG1 (50 nmol/L) with 8-oxoGua-containing DNA substrate (50 nmol/L) with a range of agonist concentrations.

| Compound | R<sub>2</sub> | R<sub>3</sub> | R<sub>4</sub> | R<sub>5</sub> | R<sub>6</sub> | R<sub>7</sub> | EC<sub>50</sub> | Δfold |
| --- | --- | --- | --- | --- | --- | --- | --- | --- |
| F02 | H | H | H | H | H | H | 2.0 ± 0.4 | 3.8 |
| F40 | phenyl | H | H | H | H | H | 1.7 ± 0.6 | 3.7 |
| F39 | pyridyl | H | H | H | H | H | 11.6 ± 1.4 | 3.7 |
| F41 | pyridyl | H | H | H | H | H | 18.2 ± 6.6 | 3.5 |
| F54 | pyridyl | H | H | H | H | H | 20.9 ± 5.9 | 2.9 |
| F52 | H | COCH<sub>3</sub> | H | H | H | H | 4.9 ± 1.3 | 3.8 |
| F20 | H | H | H |  | H | H | - | 1.1 |
| F05 | H | H | H | cyclohexyl<br>pyrimidine | H | H | - | 1.0 |
| F14 | H | H | H | imidazole | H | H | - | 1.2 |
| F23 | H | H | H | methyl benzoate | H | H | - | 0.9 |
| F10 | H | H | H | quinoline | H | H | - | 1.3 |
| F03 | H | H | H | 2-F-4,5-dimethoxyphenyl | H | H | - | 1.1 |
| F21 | H | H | H | 2-furanyl-phenyl | H | H | - | 1.1 |
| F08 | H | H | H | 2-pyrazole-5-F-phenyl | H | H | - | 1.0 |
| F07 | H | H | H | 3-Cl-4-OH-phenyl | H | H | - | 1.1 |
| F15 | H | H | H | H | 2-F-6-methoxyphenyl | phenyl | - | 1.0 |
| F16 | H | H | H | H | 3-OH-phenyl | phenyl | - | 1.0 |
| F13 | H | H | H | H | 4-CN-phenyl | phenyl | - | 1.1 |
| F18 | H | H | H | H | allyl pyrazole | phenyl | - | 1.0 |
| F24 | H | H | H | H | ethyl<br>pyrimidine | phenyl | - | 0.8 |
| F06 | H | H | H | H | ethylimidazole | phenyl | - | 1.0 |
| F11 | H | H | H | H | furan | phenyl | - | 1.0 |
| F22 | H | H | H | H | pyridyl | phenyl | - | 1.0 |

**Table 2.** R_1_ 4-OH-phenyls. The fold change of the initial reaction rate of OGG1 (50 nmol/L) with 8-oxoGua-containing DNA substrate (50 nmol/L) in the presence of agonists (10 µmol/L) is given relative to the DMSO control. EC_50_ concentrations determined by analyzing initial reaction rates of OGG1 (50 nmol/L) with 8-oxoGua-containing DNA substrate (50 nmol/L) with a range of agonist concentrations.

| Compound | R<sub>2</sub> | R<sub>3</sub> | R<sub>4</sub> | R<sub>5</sub> | R<sub>6</sub> | R<sub>7</sub> | EC<sub>50</sub> | Δfold |
| --- | --- | --- | --- | --- | --- | --- | --- | --- |
| F12 | H | H | CH<sub>3</sub> | benzofurane | H | H | - | 1.0 |
| F09 | H | H | CH<sub>3</sub> | butylimidazole | H | H | - | 1.2 |
| F19 | H | H | CH<sub>3</sub> | N-ethyl<br>pyrimidine amine | H | H | 91.7 | 1.4 |
| F04 | H | H | CH<sub>3</sub> | phenyloxazole | H | H | - | 1.4 |
| F17 | H | H | H | thiophene-<br>propynol | H | H | - | 1.2 |

**Table 3.** R_1_ halogenated phenyls. The fold change of the initial reaction rate of OGG1 (50 nmol/L) with 8-oxoGua-containing DNA substrate (50 nmol/L) in the presence of agonists (10 µmol/L) is given relative to the DMSO control. EC_50_ concentrations determined by analyzing initial reaction rates of OGG1 (50 nmol/L) with 8-oxoGua-containing DNA substrate (50 nmol/L) with a range of agonist concentrations.

| Compound | R<sub>1</sub> | R<sub>2</sub> | R<sub>3</sub> | R<sub>4</sub> | R<sub>5</sub> | R<sub>6</sub> | R<sub>7</sub> | EC<sub>50</sub> | Δfold |
| --- | --- | --- | --- | --- | --- | --- | --- | --- | --- |
| F47 | 3-Br-phenyl | H | H | H | H | H | H | 2.9 ± 0.1 | 3.5 |
| F46 | 4-Br-phenyl | H | H | H | H | H | H | 25.6 ± 10.4 | 2.2 |
| F43 | 4-Cl-phenyl | H | H | H | H | H | H | 11.0 ± 1.5 | 3.3 |
| F44 | 4-F-phenyl | H | H | H | H | H | H | 2.3 ± 1.6 | 2.5 |
| F53 | 4-F-phenyl | pyridyl | H | H | H | H | H | 20.2 ± 3.7 | 4.0 |
| F45 | 3-n-propyl<br>phenyl | H | H | H | H | H | H | - | 0.5 |

**Table 4.** R_1_ 2,4-dichloro phenyls. The fold change of the initial reaction rate of OGG1 (50 nmol/L) with 8-oxoGua-containing DNA substrate (50 nmol/L) in the presence of agonists (10 µmol/L) is given relative to the DMSO control. EC_50_ concentrations determined by analyzing initial reaction rates of OGG1 (50 nmol/L) with 8-oxoGua-containing DNA substrate (50 nmol/L) with a range of agonist concentrations.

| Compound | R<sub>2</sub> | R<sub>3</sub> | R<sub>4</sub> | R<sub>5</sub> | R<sub>6</sub> | R<sub>7</sub> | EC<sub>50</sub> | Δfold |
| --- | --- | --- | --- | --- | --- | --- | --- | --- |
| F42 | H | H | H | H | H | H | 7.5 ± 0.4 | 3.3 |
| F56 | methyl | H | H | H | H | H | - | 1.5 |
| F57 | ethyl | H | H | H | H | H | - | 1.6 |
| F58 | isopropyl | H | H | H | H | H | 23.1 ± 0.4 | 2.0 |
| F59 | cyclopentyl | H | H | H | H | H | 17.6 ± 3.5 | 2.3 |
| F01 | cyclohexyl | H | H | H | H | H | 9.1 ± 0.5 | 2.7 |
| F55 | phenyl | H | H | H | H | H | 21.2 ± 2.3 | 2.0 |
| F48 | H | COCH<sub>3</sub> | H | H | H | H | 17.4 ± 4.7 | 2.8 |
| F51 | H | benzyl | H | H | H | H | 3.8 ± 0.2 | 2.0 |
| F50 | H | 4-Cl-benzyl | H | H | H | H | 2.8 ± 0.5 | 3.1 |
| F49 | H | H | 4-dimethyl<br>amino<br>benzyl | H | H | H | 18.8 ± 5.4 | 2.4 |

**Table 5.** R_1_ CH_2_-O-phenyls. The fold change of the initial reaction rate of OGG1 (50 nmol/L) with 8-oxoGua-containing DNA substrate (50 nmol/L) in the presence of agonists (10 µmol/L) is given relative to the DMSO control. EC_50_ concentrations determined by analyzing initial reaction rates of OGG1 (50 nmol/L) with 8-oxoGua-containing DNA substrate (50 nmol/L) with a range of agonist concentrations.

| Compound | R<sub>1</sub> | R<sub>2</sub> | R<sub>3</sub> | R<sub>4</sub> | R<sub>5</sub> | R<sub>6</sub> | R<sub>7</sub> | EC<sub>50</sub> | Δfold |
| --- | --- | --- | --- | --- | --- | --- | --- | --- | --- |
| F28 | CH<sub>2</sub>-O-phenyl | H | H | H | H | H | H | 23.0 ± 6.7 | 1.6 |
| F31 | CH<sub>2</sub>-O-phenyl-4-Br | H | H | H | H | H | H | - | 1.8 |
| F36 | CH<sub>2</sub>-O-phenyl-4-Cl | H | H | H | H | H | H | - | 1.4 |
| F35 | CH<sub>2</sub>-O-phenyl-4-Cl | H | H | H | CH<sub>3</sub> | H | H | - | 1.2 |

**Table 6.** R_1_ pyridyls. The fold change of the initial reaction rate of OGG1 (50 nmol/L) with 8-oxoGua-containing DNA substrate (50 nmol/L) in the presence of agonists (10 µmol/L) is given relative to the DMSO control.

| Compound | R<sub>2</sub> | R<sub>3</sub> | R<sub>4</sub> | R<sub>5</sub> | R<sub>6</sub> | R<sub>7</sub> | EC<sub>50</sub> | Δfold |
| --- | --- | --- | --- | --- | --- | --- | --- | --- |
| F34 | H | H | H | Cl | H | H | - | 1.1 |
| F37 | H | H | H | pyridyl | H | H | - | 1.1 |

**Table 7.** Other. The fold change of the initial reaction rate of OGG1 (50 nmol/L) with 8-oxoGua-containing DNA substrate (50 nmol/L) in the presence of agonists (10 µmol/L) is given relative to the DMSO control. EC_50_ concentrations determined by analyzing initial reaction rates of OGG1 (50 nmol/L) with 8-oxoGua-containing DNA substrate (50 nmol/L) with a range of agonist concentrations.

| Compound | Structure | EC<sub>50</sub> | Δfold |
| --- | --- | --- | --- |
| F25 |  | - | 0.9 |
| F26 |  | 4.4 ± 0.3 | 4 |
| F27 |  | - | 1.1 |
| F29 |  | 15.9 ± 1.7 | 3 |
| F30 |  | - | 1.5 |
| F32 |  | - | 1.6 |
| F33 |  | - | 0.9 |
| F38 |  | 36.1 ± 3.0 | 2.0 |

### Identification of compounds that stimulate the overall kinetic rate of OGG1

Consistent with the previous report (39), independent synthesis and analyses of the activity of F01 confirmed it as an activator of OGG1 using an 8-oxoGua-containing DNA substrate, with an EC_50_ = 9.1 µmol/L ± 0.5 µmol/L (Fig. 1C) and a fold stimulation of 2.7 at 10 µmol/L (Table 4; Fig. 2B). To address our hypothesis that structural derivatives of F01 may further enhance OGG1 activity, all compounds were initially screened at 10 µmol/L for stimulation of OGG1 activity in a fluorescence-based DNA cleavage assay with a 17-mer oligodeoxynucleotide containing an 8-oxoGua at sixth position (Fig. 2A). The screen using the 8-oxoGua-containing substrate identified an additional 22 compounds that stimulated OGG1 activity by at least 2-fold relative to the DMSO control (Fig. 2B and Tables 1-7): six molecules with R_1_ phenyl substitutions, five molecules with R_1_ halogenated phenyl substitutions, eight molecules with R_1_ 2,4-dichloro phenyl substitutions (F01 is a member of this group), and 3 additional compounds that did not fit into the core structure shown in Fig. 1B. At 10 µmol/L, molecules with R_1_ 4-OH-phenyl (Table 2), R_1_ CH_2_-O-phenyl (Table 5) or pyridyl (Table 6) substitutions provided minimal or no stimulation of OGG1 activity. Additionally, compounds with any R_5_, R_6_, or R_7_ substitutions to the core molecular structure did not stimulate OGG1 activity using the 8-oxoGua-containing substrate. R_5_, R_6_, and R_7_ positions correspond to substitutions on the imidazole ring of the core molecular structure.

**Figure 2:**
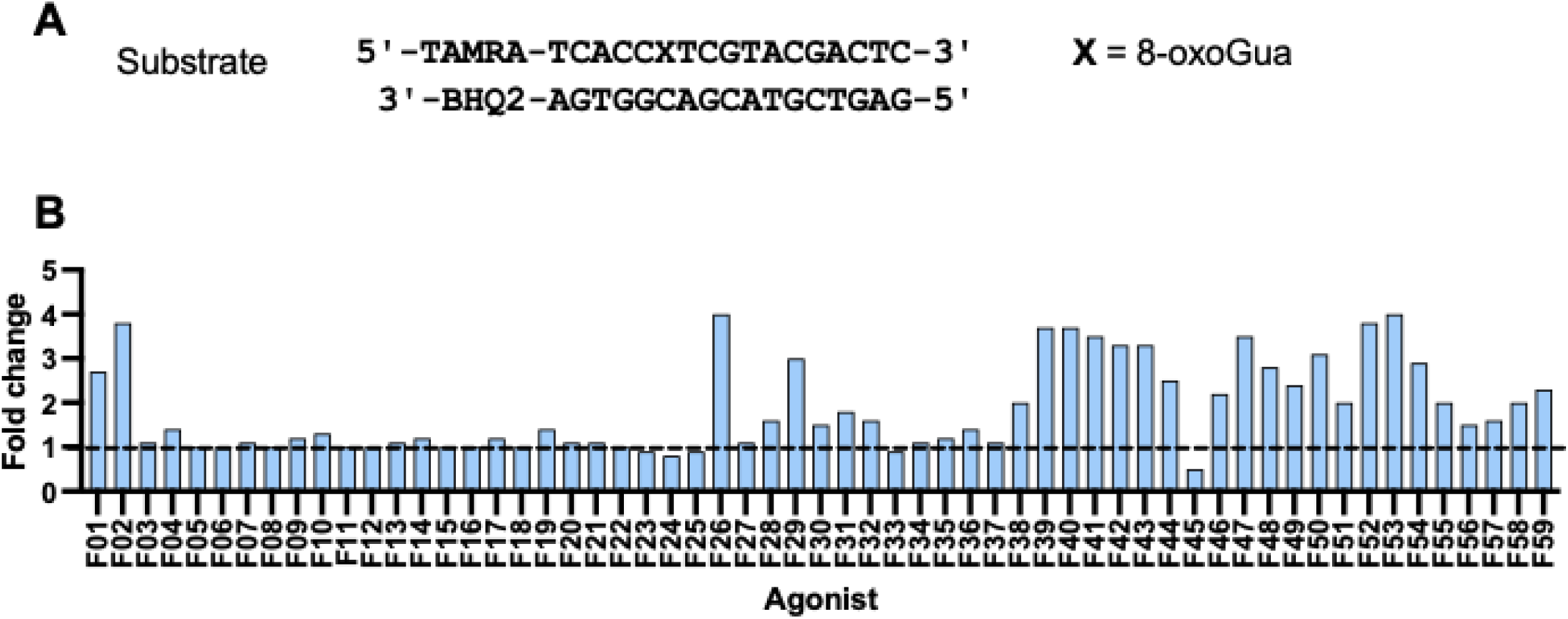
Effect of compounds on OGG1 activity with 8-oxoGua-containing substrate. *A,* 8-oxoGua-containing DNA substrate. *B*, Plot showing fold change of initial reaction rate of OGG1 (50 nmol/L) with 8-oxoGua-containing DNA substrate (50 nmol/L) in the presence of agonists (10 µmol/L) relative to the DMSO control (dashed line). Product formation was monitored by measuring TAMRA fluorescence every 2 min for 1 h. The initial rate of each reaction was determined by applying a linear fit to the linear phase of the reaction. The slope of each linear trend line was divided by the slope of the OGG1 + DMSO control reaction to calculate the relative fold change.

Based on this initial screen and using an 8-oxoGua-containing DNA substrate, the 22 compounds that stimulated OGG1 activity by ≍2-fold or greater and F19 in the R_1_ 4-OH phenyl group and F28 in the R_1_CH2-O-phenyl group were further characterized by measurements of EC_50_ concentration (Tables 1-7, Fig. S2). EC_50_ concentrations were determined using the same fluorescence-based DNA cleavage assay described above, using the 8-oxoGua-containing DNA substrate and increasing concentrations of agonist. Changes in the initial incision rates were fitted to a sigmoidal function, with the EC_50_ values calculated to be the concentration of the compound that was required to generate a 50 % increase in activity relative to the maximum stimulation. In addition to these molecules, we also independently determined the EC_50_ for the previously reported OGG1 agonist, TH10785 (Fig. S2). The EC_50_ concentrations ranged from 1.7 µmol/L ± 0.6 µmol/L (F40) to 36.1 µmol/L ± 3.0 µmol/L (F38). Nine compounds were identified with lower EC_50_ concentrations than F01 (F01: EC_50_ = 9.1 µmol/L ± 0.5 µmol/L Fig. 1C) and six were lower than or comparable to TH10785 (EC_50_ = 3.6 µmol/L ± 1.2 µmol/L (Fig. S2). Subsets of molecules with R_1_ phenyl and 2,4-dichloro were notable, with several compounds within each group having EC_50_ values less than F01. Interestingly, the EC_50_ concentration of molecules with R_1_ halogenated phenyl substitutions, specifically molecules with halogens at the C_4_ position of the phenyl moiety (F43, F44, and F46), was dependent on the type of halogen added to the phenyl, with more electronegative halogens providing lower EC_50_ concentrations (Table 3). Additionally, there was no correlation between EC_50_ concentrations, and the magnitude of fold-increase provided to OGG1 activity by compounds at 10 µmol/L (analyses not shown). Although these comparative analyses revealed the superior potency of several compounds relative to previously identified OGG1-stimulating molecules, the limitation of the assay is that it is a measure of the overall catalytic efficiency of the agonists on OGG1, but does not provide insights into which step in the reaction pathway is being accelerated, since stimulation could occur at either of the chemistry reactions or the product dissociation/enzyme turnover step.

### OGG1 agonists do not stimulate the activity of other glycosylases with overlapping substrate specificities

To investigate the specificity of these agonists for the stimulation of OGG1, fluorescence-based DNA cleavage assays were performed with human NEIL1, NTH1, and *E. coli* Fpg. Evaluating the effects of OGG1 agonists on NEIL1, NTH1, and Fpg activities was of interest because these glycosylases share substrate specificity with OGG1. NEIL1, NTH1, Fpg, and OGG1 excise Fapy lesions, while Fpg and OGG1 both excise 8-oxoGua [reviewed in (3)]. DNA cleavage assays with Fpg (15 nmol/L) and a selected group of OGG1 agonists (10 µmol/L) were performed using the 8-oxoGua-containing oligodeoxynucleotide. These agonists did not significantly stimulate Fpg activity on this substrate (Fig. S3A). DNA cleavage assays with WT edited NEIL1 (15 nmol/L) or NTH1 (11 nmol/L) and the same group of agonists (10 µmol/L) were performed using a ThyGly-containing oligodeoxynucleotide (Fig. S3B & C, respectively). These agonists did not significantly stimulate NEIL1 activity. However, F02 (10 µmol/L) completely inhibited NEIL1 activity with the ThyGly-containing DNA substrate (Fig. S3B). Inhibition of NEIL1 by F02 was found to be concentration dependent with an IC_50_ of ≍2.0 µmol/L (data not shown). The same group of agonists did not stimulate or inhibit NTH1 activity with the ThyGly substrate (Fig. S3C). The lack of stimulation of Fpg, NEIL1, and NTH1 activity by OGG1 agonists demonstrates the specificity of these molecules to enhance OGG1 activity. These data also provide evidence that these agonists do not increase the thermal denaturation temperature of oligodeoxynucleotide substrates in the fluorescence-based DNA cleavage assay.

#### OGG1 agonists increase OGG1-catalyzed release of 8-oxoGua and FapyGua from high molecular weight DNA

Prior investigations have demonstrated that the substrate range of OGG1 not only included 8-oxoGua, but also FapyGua (6, 50). To determine if these agonists also increased the rate of release of FapyGua, OGG1 ± selected agonists were reacted with γ-irradiated calf thymus DNA, and the amount of released 8-oxoGua and FapyGua bases measured by GC-MS/MS analyses. The relative abundance of these substrates following ionizing radiation is approximately equivalent (3, 51, 52). This assay directly measures the amounts of released 8-oxoGua and FapyGua and indirectly reflects the combined downstream factors that ultimately contribute to enzyme turnover. Thus, any stimulation of OGG1 activity in this assay can be accounted for by either an increased rate of the glycosylase activity of OGG1 or by an increased rate of OGG1 turnover. To ensure that the concentrations of OGG1 in the reactions were appropriate to measure increases in OGG1-catalyzed base release, γ-irradiated genomic DNA was reacted for 10 min with increasing concentrations of OGG1, thus establishing limiting enzyme conditions for the excision of 8-oxoGua (Fig. 3A) and FapyGua (Fig. 3B). OGG1 agonists were tested at concentrations 2-fold higher than the EC_50_ concentrations as previously determined using the fluorescence-based DNA cleavage assays. In reactions containing 2 µg of OGG1, significantly more 8-oxoGua was excised from DNA in the presence of F01, F02, F26, F50, and F55 relative to the DMSO control reaction (Fig. 3C). F29, F49, and F58 did not increase the amount of 8-oxoGua excised by OGG1. Under the same conditions, F01 and F26 significantly increased the amount of FapyGua excised from DNA relative to the OGG1 DMSO control (Fig. 3D). TH10785 significantly inhibited the amount of both 8-oxoGua and FapyGua excised by OGG1. Additionally, F58 significantly inhibited the amount of FapyGua excised by OGG1.

**Figure 3:**
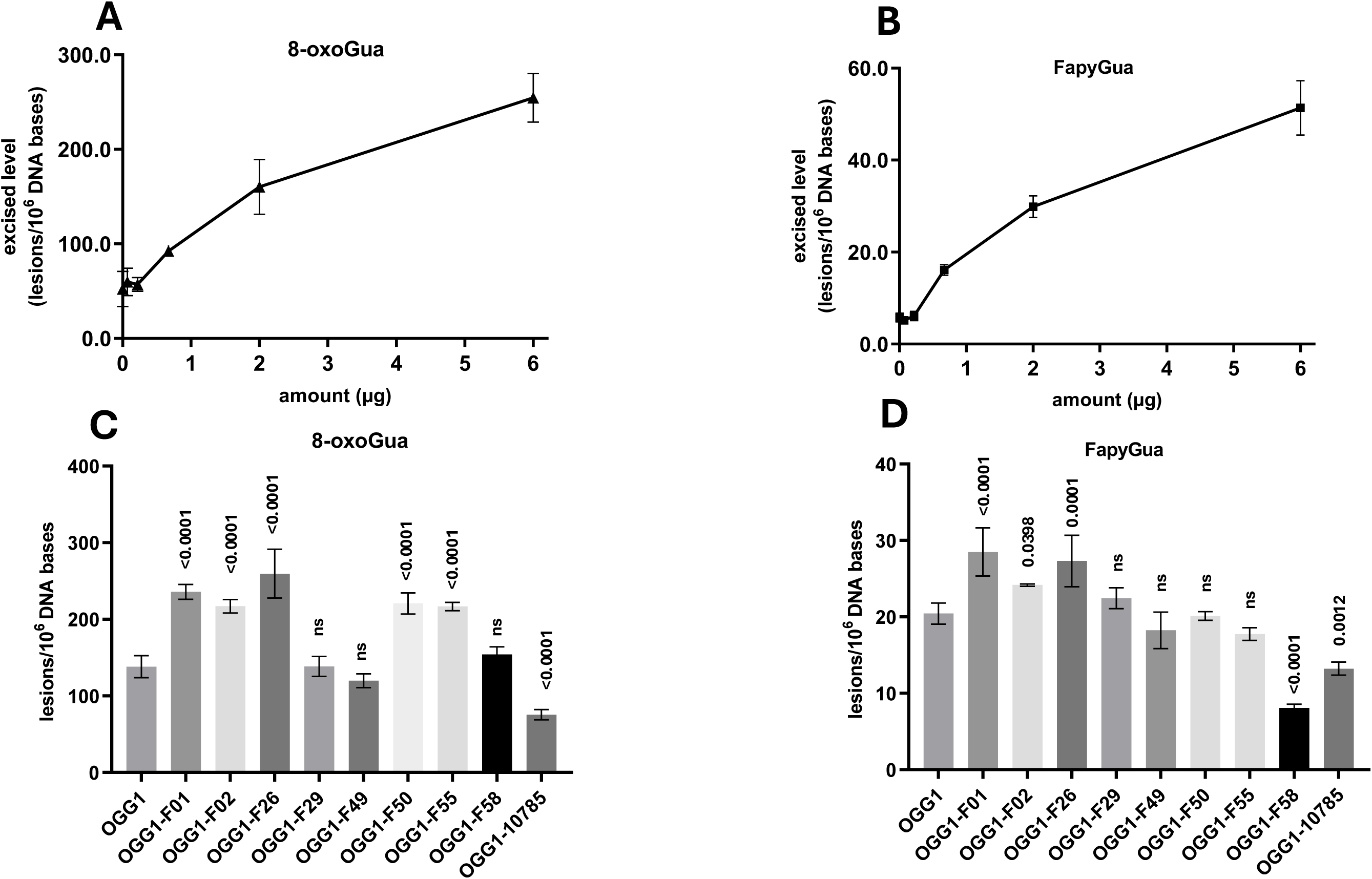
Effect of OGG1 agonists on the excision of 8-oxoGua and FapyGua from high molecular weight DNA. *A,* Levels of 8-oxoGua and *B,* FapyGua excised by increasing OGG1 concentrations from γ-irradiated calf thymus DNA were measured by GC-MS/MS at various OGG1 concentrations. *C,* Levels of 8-oxoGua and *D,* FapyGua excised by 2 µg OGG1 in the presence of agonist at 2-fold their EC_50_ concentrations from γ-irradiated calf thymus DNA were measured by GC-MS/MS at 10 min. Uncertainties are standard deviations.

### OGG1 agonists stimulate the OGG1 AP lyase reaction, but not the glycosylase reaction

Kinetic rates derived from the fluorescence-based DNA cleavage assay with 8-oxoGua-containing DNA described above reflect the combined overall rates of the glycosylase, AP lyase, enzyme dissociation, and the thermal denaturation of duplex DNA necessary to release the fluorescent signal for detection. To independently evaluate whether the agonists could stimulate the AP lyase activity of OGG1 without initiating the chemistry via a damaged base, a comparable fluorescence-based DNA cleavage assay to the one described above was performed with a 17-mer oligodeoxynucleotide containing a site-specific AP site (Fig. 4A). AP sites were generated from uracil-containing duplex DNAs by incubation with UDG, followed by reactions containing OGG1 and agonists at a 10 µmol/L final concentration. These analyses revealed that all compounds which stimulated OGG1 activity on the 8-oxoGua-containing substrate, also stimulated OGG1 activity with AP site-containing substrate (Fig. 4B). In all cases where stimulation was observed, the magnitude of OGG1 stimulation with AP site-containing DNA was either equal to or greater than the magnitude of stimulation observed with 8-oxoGua-containing DNA. To control for the possibility that OGG1 agonists alone can cleave AP site-containing DNA, 10 µmol/L agonists were incubated with the DNA substrate, and fluorescence was monitored every 2 min for 1 h. Fluorescence levels did not increase in the presence of any of the compounds, indicating that OGG1 agonists do not cleave DNA at AP sites (Fig. S4).

**Figure 4:**
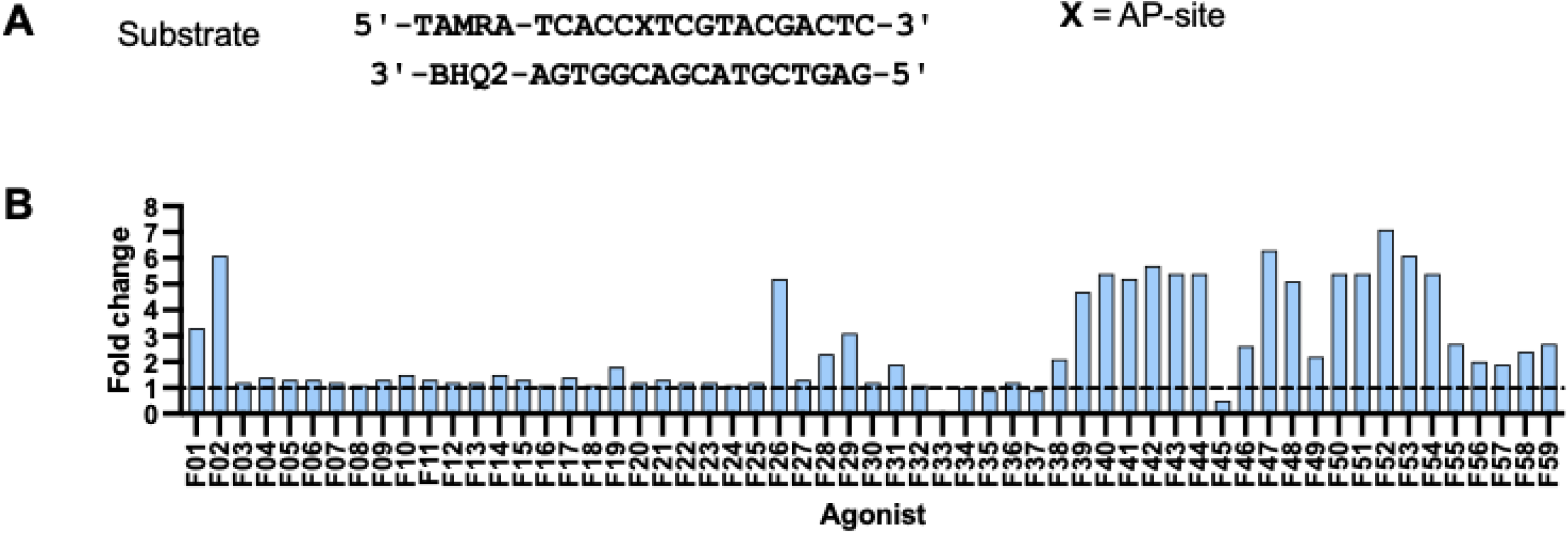
Effect of compounds on OGG1-catalyzed AP lyase activity. *A,* AP site-containing DNA substrate. *B,* Plot showing fold change of the initial reaction rate of OGG1 with AP site-containing DNA substrate in the presence of agonists (10 µmol/L) relative to the DMSO control (dashed line). Product formation was monitored by measuring TAMRA fluorescence every 2 min for 1 h. The initial rate of each reaction was determined by applying a linear fit to the linear phase of the reaction. The slope of each linear trend line was divided by the slope of the OGG1 + DMSO control reaction to calculate the relative fold change.

Although these data demonstrated agonist-mediated stimulation of OGG1 AP lyase activity, this experimental design was not informative concerning the glycosylase activity. To address this, we utilized prior studies by Dalhus et al. (53) who used site-directed mutagenesis to create separation-of-function OGG1 mutants, in which specific mutations were introduced that eliminated the AP lyase activity, while retaining the glycosylase activity. Specifically, this previous investigation demonstrated that the KCCK OGG1 double mutant retained the ability to excise 8-oxoGua from oligodeoxynucleotides, without catalyzing the AP lyase reaction (53). The KCCK mutant enzyme was purified using the same *E. coli* expression and purification procedures as that used to generate the WT OGG1.

The effects of selected agonists on the DNA glycosylase activity of the KCCK mutant were evaluated at 10 µmol/L in a gel-based DNA-cleavage assay with the glycosylase-only KCCK OGG1 mutant and 8-oxoGua-containing oligodeoxynucleotide. Since the KCCK OGG1 mutant cannot catalyze the AP lyase chemistry, sodium hydroxide was used to both hydrolyze AP sites and terminate reactions. After 15 min reactions, equivalent product formation was observed in the presence or absence of F01, F02, F26, F29, F50, F55, F58, F59 and TH10785 (Fig. S5). These data demonstrate that none of these compounds could stimulate the glycosylase activity of the KCCK OGG1 mutant with 8-oxoGua-containing DNA.

### OGG1 agonists protect against PQ-induced cytotoxicity

As part of the prior study that identified F01 as an OGG1 agonist, PQ, a formerly widely-applied herbicide, was used as a mitochondrial stressor based on its known mechanism of action (54, 55). PQ undergoes redox cycling in the mitochondrial matrix by complex I to form the PQ radical cation, reacting with oxygen to form superoxide. The production of superoxide anions in close proximity to mitochondrial DNA leads to increases in oxidatively-induced base damage (56, 57). In this prior investigation, F01 was able to rescue type II alveolar A549 epithelial cells from a variety of PQ-induced toxicities. Building off of this prior precedent and the demonstration that PQ-induced toxicity is mediated via mitochondrial DNA damage, we hypothesized that agonists identified in this investigation could mitigate the impact of PQ-induced cytotoxicity by stimulating OGG1 activity.

To evaluate this hypothesis, we first established the effects of PQ and selected OGG1 agonists on the metabolic activity of the human acute myeloid leukemia (AML) cell line, Kasumi-1, which expresses ≍8-fold lower levels of OGG1 relative to control AML cells (49). Exponentially-growing Kasumi-1 cells were treated with increasing concentrations of selected agonists ranging from (0 µmol/L to 10 µmol/L) for 72 h, with no significant decreases in metabolic activity for any compound tested (data not shown). Cells were then treated with either PQ (0 µmol/L to 500 µmol/L) (Fig. 5A) or selected OGG1 agonists (F01, F02, F49, F50, F51, F55, F58, and F59) at 10 µmol/L (Fig. 5B) and assayed for decreased metabolic activity after 72 h using an AlamarBlue assay. PQ treatment resulted in decreased metabolic activity in a dose dependent manner relative to untreated cells with a D_50_ of ≍ 200 µmol/L (Fig. 5A). For treatments with agonists alone at 10 µmol/L, the relative metabolic activities were not significantly different than that of DMSO controls (Fig. 5B).

**Figure 5:**
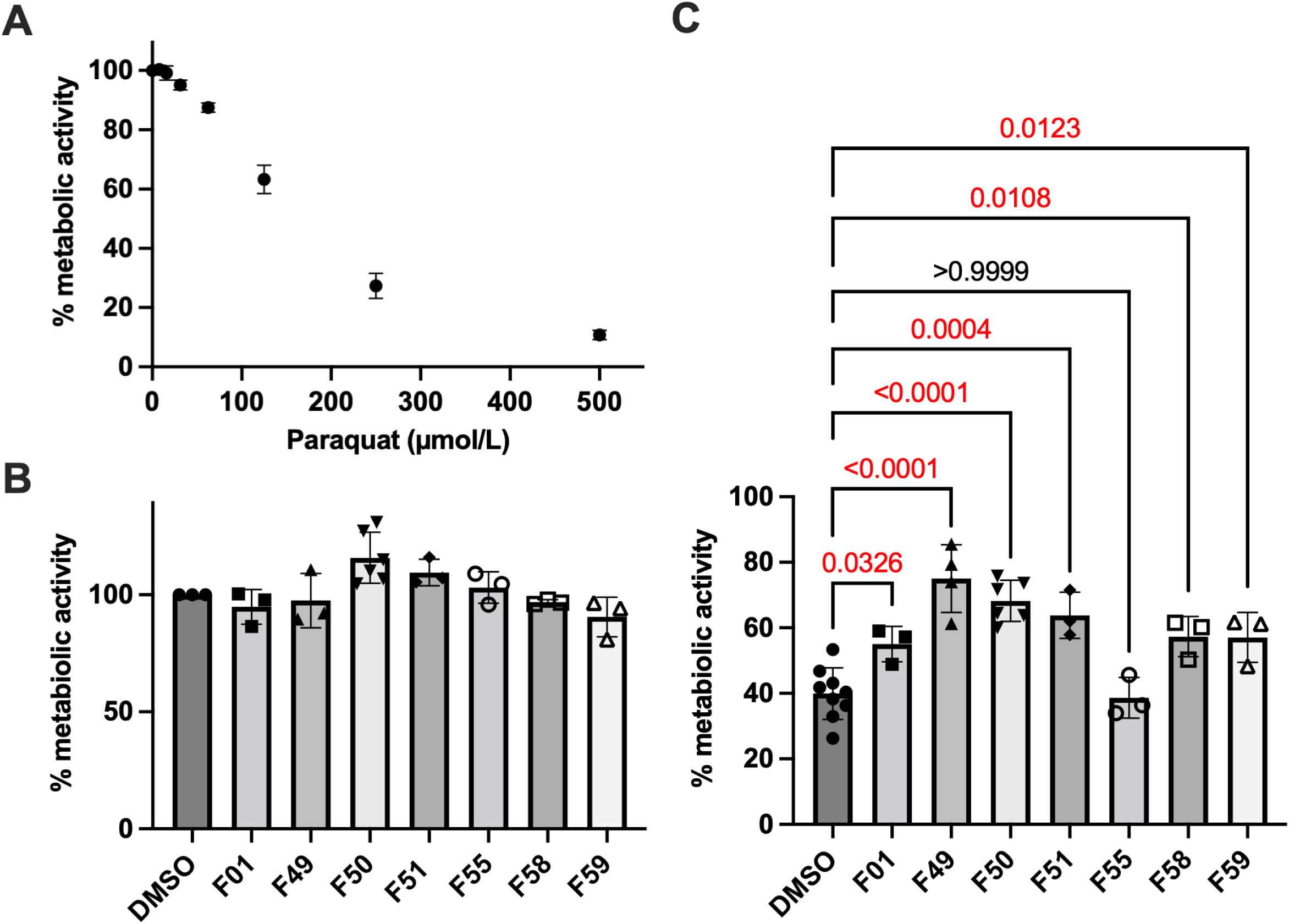
OGG1 agonists protect against PQ-induced cytotoxicity. Kasumi-1 cells were treated with increasing concentrations of either *A,* PQ, or *B*, selected agonists and the relative metabolic activity was determined. *C*, Cells were pre-treated with OGG1 agonists at the specified concentration for 3 h and then treated with 250 µmol/L PQ. Cells were incubated for 72 h after treatment and relative metabolic activity was assayed using AlamarBlue. Fluorescence readings from agonist treated cells were normalized to control cells treated with 0.1 % (v/v) DMSO. Means of three biological replicates and standard deviations are plotted with significance calculated using an ordinary one-way ANOVA test.

To test our hypothesis that agonists could protect against PQ-induced cytotoxicity, exponentially-growing Kasumi-1 cells were treated with 10 µmol/L OGG1 agonists for 3 h prior to treatment with 250 µmol/L PQ and relative metabolic activity assayed via AlamarBlue 72 h later. These data demonstrated that while F55 did not increase metabolic activity of cells treated with PQ, F01, F49, F50, F51, F58, and F59 conferred significant protection from PQ-induced decreases in metabolic activity (Fig. 5C).

## Discussion

### Enhancement of BER as a therapeutic strategy

Oxidatively-induced DNA damage from both mitochondrial metabolism and exogenous sources presents a constant challenge to cells. The efficient clearance of mutagenic 8-oxoGua and FapyGua lesions from the nuclear and mitochondrial genomes is essential for the maintenance of genomic stability. By initiating the BER pathway, OGG1 plays a fundamental role in the repair of these lesions and may affect the biological fate of cells under conditions of oxidative stress since 8-oxoGua accumulates in post-mitotic cells such as neurons in Alzheimer’s (58) and Parkinson’s disease (24), as well as in the serum of type 2 diabetes patients (59). In addition to the accumulation of oxidatively-induced DNA damage, the aforementioned disease states also share a hallmark of mitochondrial dysfunction and are correlated with OGG1-deficiency (16–20).

Of the eight human isoforms of OGG1, seven are targeted to the mitochondria, where these isoforms are the only enzyme with the ability to excise 8-oxoGua from mitochondrial DNA (60). The age- and diet-induced metabolic syndrome resulting from the murine knockout of *Ogg1* has been attributed to mitochondrial dysfunction, resulting from the lack of repair of oxidative base damage in mitochondrial DNA, consequently leading to a loss in cellular energy homeostasis (31–33). The prevention of age- and diet-induced metabolic syndrome, as well as the genetically-driven metabolic syndrome associated with the Agouti mouse model, has been demonstrated in a transgenic mouse model in which a 2-3-fold overexpression of human mtOGG1 fully corrected or prevented these diseases. These data demonstrated that OGG1-initiated mitochondrial DNA repair is essential for the maintenance of mitochondrial integrity and cellular energy homeostasis (34–36).

Herein, we have identified and characterized small molecules that activate OGG1 and mitigate PQ-induced cytotoxicity in OGG1-deficient human cells. The activity of these molecules was shown to be highly specific to OGG1 as the activity of other glycosylases with overlapping substrate specificities were unaffected by all but one compound. Several compounds identified in this investigation demonstrated improved EC_50_ concentrations, with the best being >5-fold improved over the original F01. Based on the disease consequences associated with deficiencies in OGG1, we hypothesize that next generations of these molecules or different chemotypes have the potential for therapeutic application in diseases mentioned above that are correlated with OGG1 deficiency.

The abilities of F49, F50, F51, F58, and F59 to mitigate PQ-induced cytotoxicity suggest that enhanced OGG1 activity removes PQ-induced oxidative base lesions, thus preventing cytotoxicity. PQ exerts its cytotoxic effects through superoxide anion production in the mitochondria (55) and thus, these findings suggest that F49, F50, F51, F58, and F59 are capable of enhancing mitochondrial OGG1 activity. Although nonsignificant, cells treated with F50 in the absences of exogenous challenge trended towards having increased metabolic activity relative to untreated control cells. It is possible that enhancement of OGG1-initiated DNA repair mitigates the effects of ROS arising from normal cellular respiration. However, further characterization of the cellular activity of these molecules will be necessary to address this possibility and explain the enhanced metabolic activity in the presence of F50 alone.

### Structure-activity relationships

Variation of the structure of OGG1 agonist candidates allowed for the analysis of their structure-activity relationship. Analyses of OGG1 activity in the presence of agonists at 10 µmol/L revealed that all substitutions to the imidazole ring of the core molecular scaffold eliminated OGG1 stimulation. This finding suggests that accessibility of the imidazole ring is necessary for OGG1 agonist activity. Conservation of and accessibility to the N_1_ and N_3_ positions of the imidazole ring may be necessary to facilitate acid and base chemistry between OGG1 and agonists at these positions. Determinations of EC_50_ for this set of compounds allowed us to further refine the structure-activity relationship. Analyses of R_1_ halogenated phenyls revealed that the EC_50_ value decreased with more electronegative halogens on the phenyl group. This suggests that having an electron-poor phenyl group increases affinity to OGG1 or facilitates OGG1 activity.

The 8-oxoGua product analogs 8-bromoGua and 8-aminoGua that activate OGG1 activity are reported to bind in the active site pocket of OGG1 and facilitate a more efficient β-elimination reaction at the AP site (37). Crystal structures of OGG1 with 8-oxoGua-containing DNA demonstrate binding of the released 8-oxoGua in the active site pocket (37). However, kinetic analyses of OGG1 reactions with 8-oxoGua-containing DNA revealed the presence of 9-deazaguanine-enhanced OGG1 AP-lyase activity (38). The 9-deazaguanine lacks the N9-atom previously thought to be required for catalysis of AP lyase chemistry by 8-bromoGua and 8-aminoGua (38). These analyses suggested an allosteric mechanism of activation in which product analogs bind outside the OGG1 active site pocket and produce a conformation change that facilitates AP lyase chemistry (38).

Co-crystal structures of TH10785 and OGG1, deposited by Michel et al. indicate that TH10785 binds in the active site pocket of OGG1 (40). This is facilitated by hydrogen bonding of nitrogen in the quinazolinamine motif of TH10785 to OGG1, with the secondary amine interacting at G42 and the basic nitrogen in the aromatic ring interacting at D268 (40). Similarity in the aromatic nitrogen rings between the quinazolinamine of TH10785 and the imidazole in our core molecular structure may indicate similar interactions with the active site pocket of OGG1. Co-crystallization of OGG1 with imidazole-based agonists will be necessary to determine if imidazole-based OGG1 agonists bind within the active site or (an) allosteric site(s) of OGG1. These data will facilitate the design of the next generation of molecules.

The core molecular structure of agonists identified in our analyses is similar to those of azole-based antifungal agents. F50, shown herein to be a potent activator of OGG1 activity with the ability to mitigate PQ-induced cytotoxicity, is the FDA approved antifungal agent econazole nitrate used to treat fungal skin infections. Econazole nitrate exerts its antifungal activity through inhibition of the fungal enzyme, sterol 14-α demethylase cytochrome P450 (CYP51), which is necessary for the synthesis of ergosterol, an essential building block of the fungal cell wall (61). Numerous biological activities exhibited by econozole nitrate have been characterized that go beyond its antifungal properties. Econazole nitrate has been reported to induce apoptosis in gastric cancer cells through activation of the p53 pathway (62). Additionally, econazole nitrate was able to induce cell death in leukemia cells by blocking Ca^2+^ channels (63) and cultured colon cancer cells by inducing G0/G1 cell cycle arrest (64). Interestingly, econozole nitrate has also been shown to be an inhibitor of lipopolysaccharide-inducible nitric oxide synthase and has an anti-inflammatory effect in rat aortic rings and murine macrophage cells (65). In the current study, F50 protected OGG1-deficient cells from PQ-induced cytotoxicity at concentrations that were not cytotoxic.

### The mechanism by which imidazole-based agonists enhance OGG1 activity

Data presented herein suggest that lead compounds enhance the AP lyase activity of OGG1 and thus, accelerate enzyme turnover. DNA cleavage assays with OGG1 and AP site-containing oligodeoxynucleotides clearly demonstrated that all agonist identified which accelerate the kinetics of OGG1 activity with 8-oxoGua containing DNA, also enhanced the OGG1-catalyzed AP lyase reaction. The finding that all agonists tested did not modulate the activity of the KCCK glycosylase-only OGG1 mutant provided evidence that these compounds do not accelerate the kinetics of OGG1 glycosylase chemistry. With this finding in mind, the detection of elevated levels of 8-oxoGua excised by OGG1 from γ-irradiated calf thymus DNA in the presence of agonists suggests that enhancement of AP lyase facilitates more efficient turnover of OGG1, thus allowing OGG1 to excise more 8-oxoGua from this high molecular weight DNA.

Previous studies have suggested that the BER role of OGG1 *in* vivo is limited to glycosylase chemistry, with APE1 performing DNA incision (10, 66). Our current findings that OGG1 agonists can mitigate PQ-induced cytotoxicity, which, as described above, enhance the AP lyase activity of OGG1, suggest that OGG1-catalyzed AP lyase chemistry is relevant *in vivo* following PQ challenge, and that enhancing the kinetics of this reaction is meaningful for cell survival. It is possible that the enhanced rate of AP lyase chemistry conferred by these agonists allows OGG1 to incise DNA at AP sites prior to displacement by APE1. This idea is also supported by the finding that TH10785, which allows OGG1 to catalyze both β-elimination and δ-elimination reactions at AP sites, is more toxic in combination with a polynucleotide kinase phosphatase (PNKP) small molecule inhibitor (40).

## Conclusion

We have identified and characterized new small molecule agonists of OGG1 with superior potency to those that have been identified previously. Data presented here suggest that imidazole-based OGG1 agonists enhance the kinetics of OGG1 catalyzed AP lyase and enhance enzyme turnover. While we have provided initial evidence for the activity of these molecules in a cell culture system, further studies will be necessary to characterize the activity of these molecules in cells. These future studies should consider not only the role of OGG1 in BER, but also the role of OGG1 in transcriptional regulation and signaling (67). Additionally, since several of the small molecules presented herein are similar to azole-based antifungal agents, which have been characterized in this context, this existing literature should be interrogated in the design of future cellular studies. Co-crystal structures of imidazole-based agonists and other OGG1 agonists will provide insight into many remaining questions and facilitate the design of next generation molecules. Small molecule enhancement of OGG1-mediated oxidative DNA damage repair has implications for several conditions including Alzheimer’s and Parkinson’s disease, as well as type 2 diabetes and aging.

## Supporting information

This article contains supporting information.

## Acknowledgements

Special thanks to Drs. Nathan Donley and Dmitri Rozanov for their assistance in developing the experimental framework for this investigation. We also thank Dr. Susan Wallace, formerly at the University of Vermont, for providing the vector used to express and purify NTH1. The authors would like to acknowledge the contributions of the OHSU Medicinal Chemistry Core (Research Resource ID: SCR 019048). During the preparation of this manuscript, our long-time friend and collaborator, Dr. Miral Dizdaroglu passed away. We are genuinely grateful for his scientific insights, contributions and pervasive enthusiasm for discovery.

## Disclaimer

Certain equipment, instruments, software, or materials are identified in this paper in order to specify the experimental procedure adequately. Such identification is not intended to imply recommendation or endorsement of any product or service by NIST, nor is it intended to imply that the materials or equipment identified are necessarily the best available for the purpose.

## Author contributions

Conceptualization: AKM, RSL, AN; Data curation: MML, IGM, SAM-G, NNT, PJ, HJ, JD; Formal analysis: MML, IGM, PJ, AN, MD, RSL, AKM; Funding acquisition: AKM, RSL; Writing – original draft: MML, AKM, RSL; Writing – review and editing: MML, IGM, PJ, AN, MD, RSL, AKM.

