## Supporting information for "Small molecule agonists of 8-oxoguanine DNA glycosylase, OGG1"

Contents: Methods, Table S1, Figures S1-S5

### *Methods*

#### *Synthesis of F01, F55, F56, F57, F58, and F59*

Unless otherwise stated, all chemicals and reagents were obtained from Thermo Fisher Scientific (Hampton, NH, USA) and were used as received. Analytical thin layer chromatography utilized DC Kieselgel 60$F_{254}$ precoated silica gel plates and spots were visualized under 254 nm UV light. Silica gel chromatography was performed using an Isolera Spektra automated purification system (Biotage, Uppsala, Sweden). HPLC analyses were performed using an Agilent 1260 Infinity instrument with detection at 254 nm. Mass spectrometry was performed using a Thermo Scientific Velos ion trap mass spectrometer (San Jose, CA) equipped with an electrospray ionization source operating in the positive ion mode. All compounds were at least 95 % pure as determined by HPLC. Fig. S1 shows the basic synthetic steps.

**1-(2,4-Dichlorophenyl)-2-(1*H*-imidazol-1-yl)ethan-1-one** (CAS # 46503-52-0). Synthesis of the common intermediate was as follows. A solution of imidazole (4.9 g, 72 mmol) in dichloromethane (DCM) (20 ml) was added dropwise into a solution of 2-chloro-1-(2,4-dichlorophenyl)ethenone (Combi-Blocks, San Diego (CA)) (5.36 g, 24 mmol) in DCM (10 ml). The resulting solution was stirred for 2 h at 40 °C and 18 h at room temperature. The reaction was concentrated, redissolved in ethyl acetate (EtOAc) (200 ml), washed three times with water (50 mL each) and twice with a saturated solution of NaCl (50 ml). The organic layer was then concentrated, redissolved in hot methanol and recrystallized overnight. The resulting product was collected by filtration and dried to give 2.91 g (47% yield) of CAS# 46503-52-0.

**General procedure for the synthesis of F01 and F55-F59.** Molecular sieves (3 Å) (200 mg) were added to a round-bottom flask and flame-dried under vacuum. After cooling, 1-(2,4-dichlorophenyl)-2-(1*H*-imidazol-1-yl)ethan-1-one (0.1 g, 0.39 mmol) was added, and the flask was sealed, evacuated, and flushed with argon. Dry tetrahydrofuran (THF) (4 mL) was added and degassed, and the solution was cooled to -78 °C in a dry ice / isopropyl alcohol bath. An appropriate Grignard reagent (RMgBr) (Sigma-Aldrich Chemical Company in St. Louis, MO (USA)) (1.6 mL, 4 mmol) was added dropwise, and the reaction was stirred in a 0 °C ice bath for 2 h. The reaction was quenched by dropwise addition of saturated aqueous ammonium chloride (10 ml). The aqueous layer was extracted three times with EtOAc (10 mL each). The organic fractions were combined, washed twice with a saturated solution of NaCl (5 mL), dried with magnesium sulfate, filtered, and concentrated. The crude material was redissolved in diethyl ether, and a solution of hydrochloric acid in dioxane was added. Recrystallization (18 h at -20 °C) resulted in yellow-orange crystals that were collected by filtration and dried to give the product as an HCl salt in approximately 10 % yield.

**1-cyclohexyl-1-(2,4-dichlorophenyl)-2-(1H-imidazol-1-yl)ethan-1-ol (F01).** F01 was prepared according to the general procedure provided above. MS for $C_{17}H_{20}Cl_2N_2O$: calculated m/z = 338.10, observed m/z = 338.10.

**1-(2,4-dichlorophenyl)-2-(1H-imidazol-1-yl)-1-phenylethan-1-ol (F55).** F55 was prepared according to the general procedure provided above. MS for $C_{17}H_{14}Cl_2N_2O$: calculated m/z = 332.05, observed m/z = 332.05.

**2-(2,4-dichlorophenyl)-1-(1H-imidazol-1-yl)propan-2-ol (F56).** F56 was prepared according to the general procedure provided above. MS for $C_{12}H_{12}Cl_2N_2O$: calculated m/z = 270.03, observed m/z = 270.03.

**2-(2,4-dichlorophenyl)-1-(1H-imidazol-1-yl)butan-2-ol (F57).** F57 was prepared according to the general procedure provided above. MS for $C_{13}H_{14}Cl_2N_2O$: calculated m/z = 284.05, observed m/z = 284.05.

**2-(2,4-dichlorophenyl)-1-(1H-imidazol-1-yl)-3-methylbutan-2-ol (F58).** F58 was prepared according to the general procedure provided above. MS for $C_{14}H_{16}Cl_2N_2O$: calculated m/z = 298.06, observed m/z = 298.06.

**1-cyclopentyl-1-(2,4-dichlorophenyl)-2-(1H-imidazol-1-yl)ethan-1-ol (F59).** F50 was prepared according to the general procedure provided above. MS for $C_{16}H_{18}Cl_2N_2O$: calculated m/z = 324.08, observed m/z = 324.08.

**Tables & Figures**

**Table S1: Source ID, name, molecular formula and molecular weight of compounds.**

| Number | Source ID | Mol Name | Mol Formula | Mol Weight |
|---|---|---|---|---|
| **F01** | OHSU MedChem Core | 1-cyclohexyl- 1-(2,4-dichlorophenyl)-2-(1H-imidazole-1-yl) ethan-1-ol | $C_{17}$ $H_{20}$ $Cl_2$ $N_2$ O | 339 |
| **F02** | ChemBridge 8881667 | 2-(1H-imidazol-1-yl)-1-phenylethan-1-ol | $C_{11}$ $H_{12}$ $N_2$ O | 188 |
| **F03** | ChemBridge 10406194 | 2-[2-(2-fluoro-4,5-dimethoxyphenyl)-1H-imidazol-1-yl]-1-phenylethan-1-ol | $C_{19}$ $H_{19}$ F $N_2$ $O_3$ | 342.4 |
| **F04** | ChemBridge 15069650 | 4-{(1S*,2R*)-1-hydroxy-2-[2-(2-phenyl-1,3-oxazol-4-yl)-1H-imidazol-1-yl]propyl}phenol | $C_{21}$ $H_{19}$ $N_3$ O3 | 361.4 |
| **F05** | ChemBridge 18775397 | 2-[2-(2-cyclohexylpyrimidin-5-yl)-1H-imidazol-1-yl]-1-phenylethan-1-ol | $C_{21}$ $H_{24}$ $N_4$ O | 348.4 |
| **F06** | ChemBridge 19083028 | 2-(1-ethyl-5'-phenyl-1H,3'H-2,4'-biimidazol-3'-yl)-1-phenylethan-1-ol | $C_{22}$ $H_{22}$ $N_4$ O | 358.4 |
| **F07** | ChemBridge 24145023 | 2-chloro-4-[1-(2-hydroxy-2-phenylethyl)-1H-imidazol-2-yl]phenol | $C_{17}$ $H_{15}$ Cl $N_2$ $O_2$ | 314.8 |
| **F08** | ChemBridge 29204921 | 2-{2-[5-fluoro-2-(1H-pyrazol-1-yl)phenyl]-1H-imidazol-1-yl}-1-phenylethan-1-ol | $C_{20}$ $H_{17}$ F $N_4$ O | 348.4 |
| **F09** | ChemBridge 29247199 | 4-[(1S*,2R*)-2-(2'-butyl-1H,1'H-2,4'-biimidazol-1-yl)-1-hydroxypropyl]phenol | $C_{19}$ $H_{24}$ $N_4$ $O_2$ | 340.4 |
| **F10** | ChemBridge 35965465 | 1-phenyl-2-(2-quinolin-2-yl-1H-imidazol-1-yl)ethan-1-ol | $C_{20}$ $H_{17}$ $N_3$ O | 315.4 |
| **F11** | ChemBridge 39135583 | 2-[5-(2-furyl)-4-phenyl-1H-imidazol-1-yl]-1-phenylethan-1-ol | $C_{21}$ $H_{18}$ $N_2$ $O_2$ | 330.4 |
| **F12** | ChemBridge 39951269 | 4-{(1S*,2R*)-2-[2-(1-benzofuran-2-yl)-1H-imidazol-1-yl]-1-hydroxypropyl}phenol | $C_{20}$ $H_{18}$ $N_2$ $O_3$ | 334.4 |
| **F13** | ChemBridge 45359558 | 4-[1-(2-hydroxy-2-phenylethyl)-4-phenyl-1H-imidazol-5-yl]benzonitrile | $C_{24}$ $H_{19}$ $N_3$ O | 365.4 |
| **F14** | ChemBridge 46577356 | 2-(1H,1'H-2,2'-biimidazol-1-yl)-1-phenylethan-1-ol | $C_{14}$ $H_{14}$ $N_4$ O | 254.3 |

| | | | | |
|---|---|---|---|---|
| **F15** | ChemBridge 55298977 | 2-[5-(2-fluoro-6-methoxyphenyl)-4-phenyl-1H-imidazol-1-yl]-1-phenylethan-1-ol | $C_{24} H_{21} F N_2 O_2$ | 388.4 |
| **F16** | ChemBridge 58417378 | 3-[1-(2-hydroxy-2-phenylethyl)-4-phenyl-1H-imidazol-5-yl]phenol | $C_{23} H_{20} N_2 O_2$ | 356.4 |
| **F17** | ChemBridge 63503268 | 4-(1-hydroxy-2-{2-[5-(3-hydroxyprop-1-yn-1-yl)-2-thienyl]-1H-imidazol-1-yl}ethyl)phenol | $C_{18} H_{16} N_2 O_3 S$ | 340.4 |
| **F18** | ChemBridge 67366050 | 2-[5-(1-allyl-1H-pyrazol-4-yl)-4-phenyl-1H-imidazol-1-yl]-1-phenylethan-1-ol | $C_{23} H_{22} N_4 O$ | 370.4 |
| **F19** | ChemBridge 71346720 | 4-((1S*,2R*)-2-{2-[2-(ethylamino)pyrimidin-5-yl]-1H-imidazol-1-yl}-1-hydroxypropyl)phenol | $C_{18} H_{21} N_5 O_2$ | 339.4 |
| **F20** | ChemBridge 78415552 | 1-phenyl-2-[2-(5,6,7,8-tetrahydro-4H-pyrazolo[1,5-a][1,4]diazepin-2-yl)-1H-imidazol-1-yl]ethan-1-ol | $C_{18} H_{21} N_5 O$ | 323.4 |
| **F21** | ChemBridge 78591809 | 2-{2-[2-(2-furyl)phenyl]-1H-imidazol-1-yl}-1-phenylethan-1-ol | $C_{21} H_{18} N_2 O_2$ | 330.4 |
| **F22** | ChemBridge 80374519 | 1-phenyl-2-(4-phenyl-5-pyridin-4-yl-1H-imidazol-1-yl)ethan-1-ol | $C_{22} H_{19} N_3 O$ | 341.4 |
| **F23** | ChemBridge 81922488 | methyl 3-[1-(2-hydroxy-2-phenylethyl)-1H-imidazol-2-yl]benzoate | $C_{19} H_{18} N_2 O_3$ | 322.4 |
| **F24** | ChemBridge 88441914 | 2-[5-(2-ethylpyrimidin-4-yl)-4-phenyl-1H-imidazol-1-yl]-1-phenylethan-1-ol | $C_{23} H_{22} N_4 O$ | 370.4 |
| **F25** | ChemBridge 4001464 | (4-fluorophenyl)(1-methyl-1H-imidazol-2-yl)methanol | $C_{11} H_{11} F N_2 O$ | 206.2 |
| **F26** | ChemBridge 4002262 | [2-(1H-imidazol-1-yl)-1-phenylethyl]amine dihydrochloride hydrate | $C_{11} H_{13} N_3 . 2 Cl H . H_2 O$ | 278.2 |
| **F27** | ChemBridge 4011409 | [1-(1H-imidazol-1-ylmethyl)propyl]amine dihydrochloride | $C_7 H_{13} N_3 . 2 Cl H$ | 212.1 |
| **F28** | ChemBridge 5107296 | 1-(1H-imidazol-1-yl)-3-phenoxy-2-propanol dihydrochloride | $C_{12} H_{14} N_2 O_2 . 2 Cl H$ | 291.2 |
| **F29** | ChemBridge 5308482 | 1-(2,4-dichlorophenyl)-2-(1H-imidazol-1-yl)ethan-1-one | $C_{11} H_8 Cl_2 N_2 O$ | 255.1 |
| **F30** | ChemBridge 5662854 | (1-methyl-1H-imidazol-2-yl)(phenyl)methanol | $C_{11} H_{12} N_2 O$ | 188.2 |

| | | | | |
|---|---|---|---|---|
| **F31** | ChemBridge 7775286 | 1-(4-bromophenoxy)-3-(1H-imidazol-1-yl)propan-2-ol | $C_{12} H_{13} Br N_2 O_2$ | 297.1 |
| **F32** | ChemBridge 8886246 | 1-phenyl-2-(1H-pyrrol-1-yl)ethan-1-one | $C_{12} H_{11} NO$ | 182.2 |
| **F33** | ChemBridge 4400175 | 1H-imidazol-1-ylacetic acid | $C_5 H_6 N_2 O_2$ | 126.1 |
| **F34** | ChemBridge 5209187 | 2-(2-chloro-1H-imidazol-1-yl)-1-(4-pyridinyl)ethan-1-ol | $C_{10} H_{10} Cl N_3 O$ | 223.7 |
| **F35** | ChemBridge 6860860 | 1-(4-chlorophenoxy)-3-(2-methyl-1H-imidazol-1-yl)propan-2-ol | $C_{13} H_{15} Cl N_2 O_2$ | 266.7 |
| **F36** | ChemBridge 6869967 | 1-(4-chlorophenoxy)-3-(1H-imidazol-1-yl)propan-2-ol | $C_{12} H_{13} Cl N_2 O_2$ | 252.7 |
| **F37** | ChemBridge 28162240 | 1-pyridin-3-yl-2-(2-pyridin-4-yl-1H-imidazol-1-yl)ethan-1-ol | $C_{15} H_{14} N_4 O$ | 266.3 |
| **F38** | ChemBridge 65017993 | N-[2-(1H-imidazol-1-yl)-1-phenylethyl]-2-methoxyacetamide | $C_{14} H_{17} N_3 O_2$ | 259.3 |
| **F39** | MolPort-002-740-873 | 2-(1H-imidazol-1-yl)-1-phenyl-1-(pyridin-3-yl)ethan-1-ol | $C_{16} H_{15} N_3 O$ | 265.316 |
| **F40** | MolPort-003-003-778 | 2-(1H-imidazol-1-yl)-1,1-diphenylethan-1-ol | $C_{17} H_{16} N_2 O$ | 264.328 |
| **F41** | MolPort-003-003-779 | 2-(1H-imidazol-1-yl)-1-phenyl-1-(pyridin-4-yl)ethan-1-ol | $C_{16} H_{15} N_3 O$ | 265.316 |
| **F42** | MolPort-001-812-137 | 1-(2,4-dichlorophenyl)-2-(1H-imidazol-1-yl)ethan-1-ol | $C_{11} H_{10} Cl_2 N_2 O$ | 257.11 |
| **F43** | MolPort-006-124-253 | 1-(4-chlorophenyl)-2-(1H-imidazol-1-yl)ethan-1-ol | $C_{11} H_{11} Cl N_2 O$ | 222.67 |
| **F44** | MolPort-008-651-069 | 1-(4-fluorophenyl)-2-(1H-imidazol-1-yl)ethan-1-ol | $C_{11} H_{11} F N_2 O$ | 206.22 |
| **F45** | MolPort-008-662-031 | 2-(1H-imidazol-1-yl)-1-(4-propylphenyl)ethan-1-ol | $C_{14} H_{18} N_2 O$ | 230.311 |
| **F46** | MolPort-008-686-825 | 1-(4-bromophenyl)-2-(1H-imidazol-1-yl)ethan-1-ol | $C_{11} H_{11} Br N_2 O$ | 267.126 |
| **F47** | MolPort-023-144-557 | 1-(3-bromophenyl)-2-(1H-imidazol-1-yl)ethan-1-ol | $C_{11} H_{11} Br N_2 O$ | 267.126 |
| **F48** | MolPort-046-532-488 | 1-(2,4-dichlorophenyl)-2-(1H-imidazol-1-yl)ethyl acetate | $C_{13} H_{12} Cl_2 N_2 O_2$ | 299.15 |
| **F49** | MolPort-008-348-956 | 1-(2,4-dichlorophenyl)-3-[4-(dimethylamino)phenyl]-2-(1H-imidazol-1-yl)propan-1-ol | $C_{20} H_{21} Cl_2 N_3 O$ | 390.31 |
| **F50** | MolPort-003-666-381 | nitric acid—1-{2-(4-chlorophenyl)-2-[(2,4-dichlorophenyl)methoxy]ethyl}-1H-imidazole (1/1) | $C_{18} H_{16} Cl_3 N_3 O_4$ | 444.69 |

| | | | | |
|---|---|---|---|---|
| **F51** | MolPort-046-691-919 | 1-[2-(benzyloxy)-2-(2,4-dichlorophenyl)ethyl]-1H-imidazole | $C_{18} H_{16} Cl_2 N_2 O$ | 347.24 |
| **F52** | MolPort-002-718-963 | 2-(1H-imidazol-1-yl)-1-phenylethyl acetate | $C_{13} H_{14} N_2 O_2$ | 230.267 |
| **F53** | MolPort-002-740-872 | 1-(4-fluorophenyl)-2-(1H-imidazol-1-yl)-1-(pyridin-2-yl)ethan-1-ol | $C_{16} H_{14} F N_3 O$ | 283.306 |
| **F54** | MolPort-002-740-917 | 2-(1H-imidazol-1-yl)-1-phenyl-1-(pyridin-2-yl)ethan-1-ol | $C_{16} H_{15} N_3 O$ | 265.316 |
| **F55** | OHSU MedChem Core | 1-(2,4-dichlorophenyl)-2-(1H-imidazol-1-yl)-1-phenylethan-1-ol | $C_{17} H_{14} Cl_2 H_2 O$ | 333.212 |
| **F56** | OHSU MedChem Core | 2-(2,4-dichlorophenyl)-1-(1H-imidazol-1-yl)propan-2-ol | $C_{12} H_{12} Cl_2 N_2 O$ | 271.141 |
| **F57** | OHSU MedChem Core | 2-(2,4-dichlorophenyl)-1-(1H-imidazol-1-yl)butan-2-ol | $C_{13} H_{14} Cl_2 N_2 O$ | 285.168 |
| **F58** | OHSU MedChem Core | 2-(2,4-dichlorophenyl)-1-(1H-imidazol-1-yl)-3-methylbutan-2-ol | $C_{14} O_{16} Cl_2 N_2 O$ | 299.195 |
| **F59** | OHSU MedChem Core | 1-cyclopentyl-1-(2,4-dichlorophenyl)-2-(1H-imidazol-1-yl)ethan-1-ol | $C_{16} H_{18} Cl_2 N_2 O$ | 325.233 |

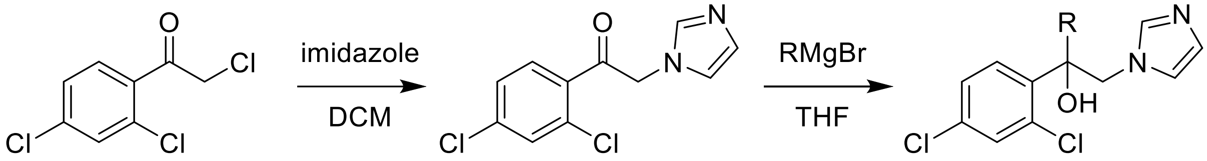


[4252-78-2]

[46503-52-0]

**F01**, R = cyclohexyl
**F55**, R = phenyl
**F56**, R = methyl
**F57**, R = ethyl
**F58**, R = isopropyl
**F59**, R = cyclopentyl

**Figure S1: Synthesis of F01, F55, F56, F57, F58 and F59.** Scheme of reaction of 2-bromo-1-(2,4-dichlorophenyl)ethenone (CAS# 4252-78-2) with imidazole and DCM, followed by the reaction of 1-(2,4-dichlorophenyl)-2-(1H-imidazol-1- yl)ethenone (CAS# 46503-52-0) with THF and alkyl magnesium bromide to provide final compounds F01, F55, F56, F57, F58, and F59.

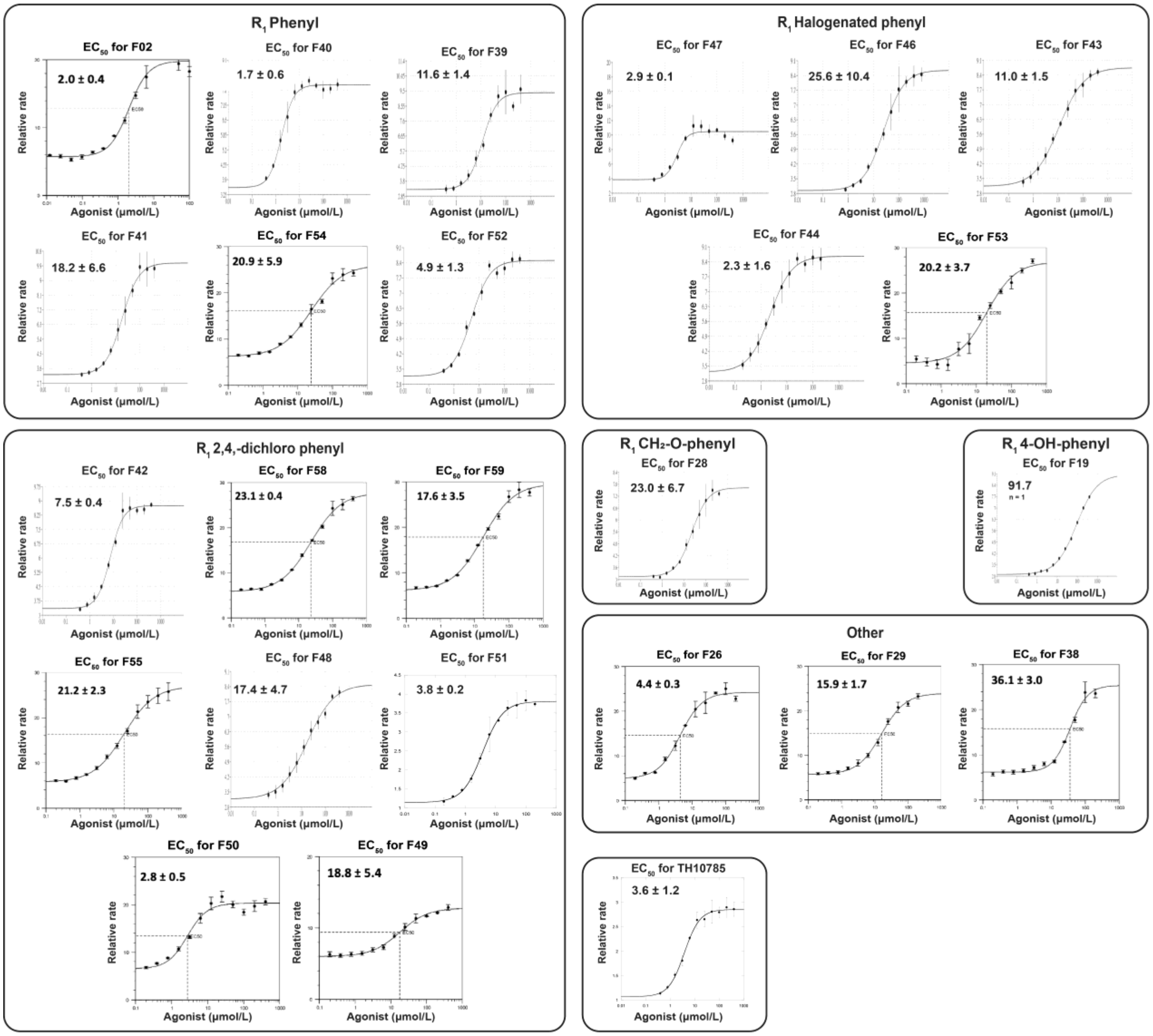


**Figure S2: $EC_{50}$ plots for agonists that stimulated OGG1 activity by 2-fold or greater in the initial screen with 8-oxoGua-containing substrate**. $EC_{50}$ concentrations were determined by analyzing relative initial reaction rates of OGG1 (50 nmol/L) on 8-oxoGua-containing DNA substrate (50 nmol/L) with a range of agonist concentrations in a fluorescence-based DNA cleavage assay. The slopes of linear trendlines at each agonist concentration were plotted, and a non-linear regression was performed. Data reflects three independent experiments with standard deviation, except for F19, which reflects a single experiment. Compounds are grouped by chemotype (see Tables 1 – 7).

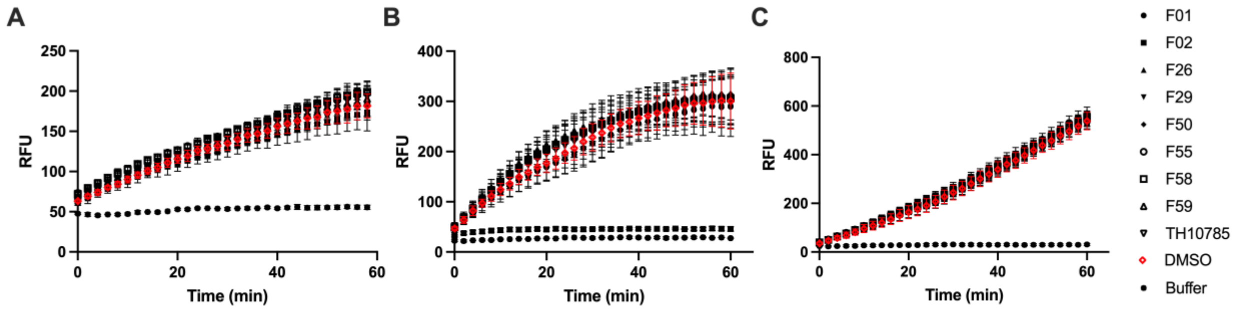


**Figure S3: Specificity of agonists to OGG1.** *A,* 8-oxoGua-containing DNA substrate was used to screen selected compounds (10 μmol/L) for stimulation of *E. coli* Fpg activity in a fluorescence-based DNA cleavage assay. *B,* ThyGly-containing DNA substrate was used to screen compounds (10 μmol/L) for stimulation of NEIL1. *C,* ThyGly-containing DNA substrate was used to screen compounds (10 μmol/L) for stimulation of NTH1 activity. Fluorescence was monitored every 2 min for 1 h using a TECAN plate reader. Uncertainties are standard deviations.

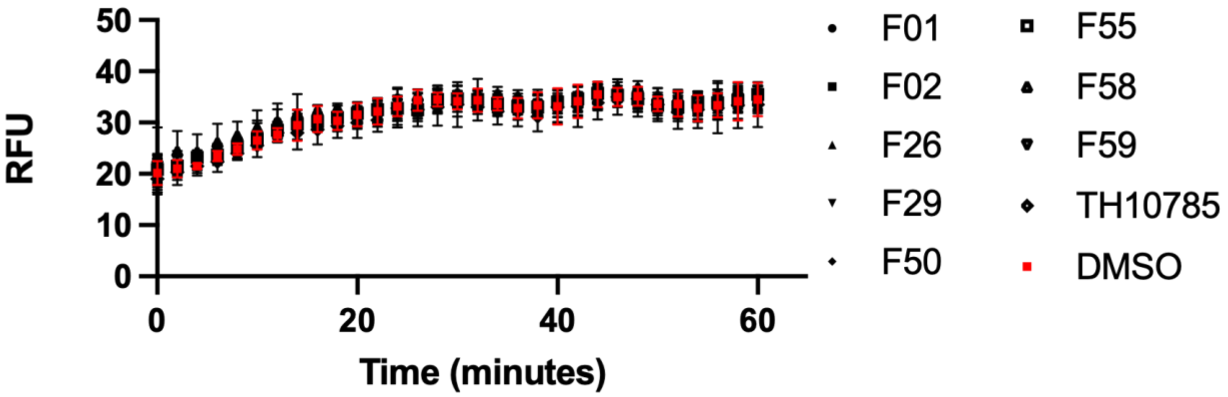


**Figure S4: Agonists alone do not cleave AP sites.** OGG1 agonists (10 µmol/L) were incubated with AP site-containing substrate (50 nmol/L) for 1 h at 37 °C. Product formation was monitored by taking fluorescence readings every 2 min for 1 h using a TECAN plate reader**.** Uncertainties are standard deviations.

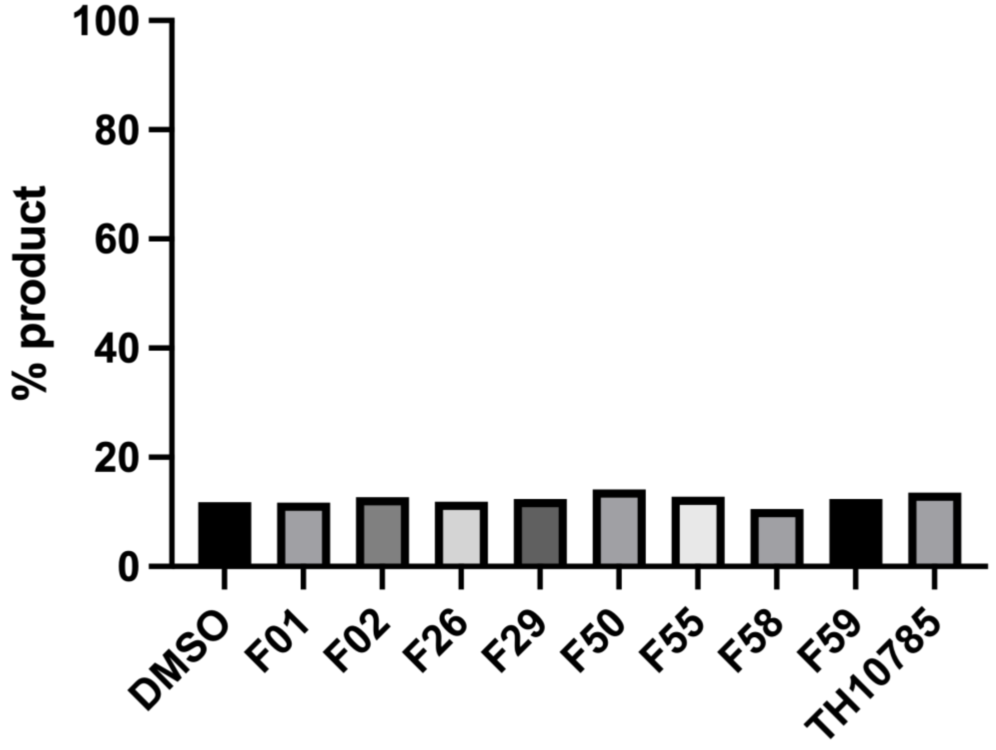


**Figure S5: OGG1 agonists do not stimulate the KCCK OGG1 mutant.** The KCCK OGG1 mutant (25 nmol/L) was reacted with 8-oxoGua-containing DNA substrate (100 nmol/L) for 10 min in the presence of 10 μmol/L agonists or DMSO. The AP sites were converted into DNA single-stranded breaks by treatment with sodium hydroxide. Products were separated by electrophoresis through a 15 % polyacrylamide gel in the presence of 8 mol/L urea and the percent product formation was determined by band density analysis and plotted.
